# ER stress drives ER-to-Golgi trafficking of ATF6 by blocking its membrane insertion

**DOI:** 10.1101/822965

**Authors:** Jia Xu, Xianbin Meng, Fang Wu, Haiteng Deng, Suneng Fu

## Abstract

Regulated ER-to-Golgi trafficking is a fundamental cellular process that enables ER-resident transcription factors to sense perturbations in the ER environment and activates transcriptional programs that restore ER and cellular homeostasis. Current models suggest sensor activation is initiated by dissociation from its ER-resident binding partners. Here we challenge this model by demonstrating that the unfolded protein sensor ATF6 is sorted to the Golgi as a newly synthesized peripheral membrane protein beyond the reach of its ER retainer, and translocon inhibition alone is sufficient to drive ATF6 activation. We identify ATF6 transmembrane domain and its C-terminus as the intrinsic factors that control membrane insertion efficiency and stress sensing capacity, and the BAG6 complex as the receptor that triages ATF6 between membrane insertion and Golgi sorting. Besides ATF6, we show that translocon inhibition expedites the activation of the cholesterol sensor SREBP2, and the catalytic domain of Golgi-bound S1P that processes both ATF6 and SREBP2 resides in the cytosol. Therefore, we propose an alternative, sensing-by-synthesis model, in which transcription factors are continuously synthesized, and perturbations to the ER environment are quantitatively sensed by the fraction of sensors that fail to be properly incorporated into ER membrane and sorted to Golgi for activation.

---

Regulated intramembrane proteolysis is an evolutionarily conserved mechanism that enables a cellular response to environmental and developmental cues ^1–3^. The accessibility of substrate protein to its protease may be regulated on multiple levels, including post-translational modification, ligand binding, and trafficking ^4,5^. Among them, how environmental and developmental signals enable the sorting of substrate protein to the protease compartment is not yet to be fully established.

The unfolded protein response (UPR) is a coordinated transcriptional and translational programs activated in response to ER stress, and they act together to reduce protein load into the ER, upregulate ER-associated protein degradation, induce chaperone expression, and eventually to restore protein homeostasis in the ER lumen ^6–8^. The induction of UPR is governed by three ER-resident, single-pass transmembrane domain proteins, namely IRE1α (inositol-requiring protein 1α), PERK (PRKR-like endoplasmic reticulum kinase), and ATF6 (activating transcription factor 6) ^9,10^.

Among the three branches of UPR, IRE1α is most conserved, and its mechanism of activation has been studied most extensively ^9^. Overwhelming data suggest that IRE1α can monitor misfolded protein accumulation in the ER lumen. First, the crystal structure of IRE1α harbors an unfolded protein binding domain ^11,12^, and binding of IRE1α by unfolded proteins induces its oligomerization and activation ^13–15^. Second, unfolded proteins may further facilitate IRE1α oligomerization by releasing it from heteromeric interaction with Grp78 ^16,17^. Additional mechanisms of IRE1α activation, although not necessarily mutually exclusive, have been reported. Dnajb9, a J-class cochaperone, has been shown to actively recruit Grp78 to monomerize and inhibit IRE1α under non-stress conditions ^18^. ER membrane lipid aberrancies may also activate IRE1α, and such activations are independent of its luminal domain ^19–23^. Even cytosolic factors, as well as chemical ligands, have been identified to be able to bind and regulate the activation of IRE1α ^24–27^.

ATF6 is a recent addition to the UPR during evolution, and it is activated through ER stress-induced ER-to-Golgi trafficking followed by proteolytic cleavage and nucleus translocation ^9^. Early work suggests that ATF6 is synthesized as a membrane-anchored transcription factor bound by Grp78 under non-stress conditions ^28^. Upon ER stress, ATF6 is released from Grp78, trafficked to the Golgi, processed by the Site-1 (S1P) and Site-2 (S2P) proteases to release its N-terminal fragment, which is then transported into the nucleus to turn on gene expression ^28–34^. A series of mutational analysis has led to the identification of Grp78 binding domains and Golgi-localization signals within the luminal domain of ATF6, and they are sufficient for ER stress sensing and Golgi targeting ^28,34,35^. However, it has been contended that the Grp78-ATF6 interaction is very stable and ATF6 is an ERAD substrate and acutely degraded upon the induction of ER stress ^36–38^. Besides, the population of ATF6 trafficked to Golgi seems to be underglycosylated and devoid of disulfide bonds ^39,40^, which may suggest that the Golgi-bound ATF6 is newly synthesized upon the initiation of ER stress. Therefore, we designed a series of experiments to dissect the exact mechanism governing the Golgi sorting of ATF6, and more broadly, the regulated trafficking of ER-tethered transcription factors.

## Results

### Golgi-bound ATF6 is unglycosylated, not deglycosylated

Unlike PERK and IRE1, both of which oligomerize in response to ER stress, ATF6 is presumed to be monomerized and deglycosylated before its transport to the Golgi for activation (Hong et al., 2004b; Nadanaka et al., 2007; Sundaram et al., 2018). However, the rapid degradation of ATF6 upon the initiation of ER stress ^37,39,40^ calls into the question about the functional utility of pre-existing ATF6 pool. Alternatively, we may hypothesize that the Golgi-targeted ATF6 is newly synthesized and therefore no need for monomerization and deglycosylation. As deglycosylation will result in the conversion of Asparagine (N) residues to Aspartate (D), we decided to test this hypothesis by examining the status of Asparagine residues in the luminal domain (C-terminus) of ATF6.

As the endogenous ATF6 protein level is low and its transition through Golgi is rapid, we expressed recombinant ATF6 protein with the S1/2P site mutated to enable its retention and visualization on the Golgi under ER stress conditions (Fig. 1a). As expected, the recombinant ATF6 was co-localized with the ER marker HSPA5 under normal conditions and sorted to GALNT2-labeled Golgi compartment upon thapsigargin (TG) treatment (Fig. 1b). Fractionation of ER and Golgi vesicles by sucrose gradient revealed an exclusive appearance of a faster-migrating and presumably non-glycosylated form of ATF6 in the Golgi fractions (Fig. 1c). The proper behavior of the recombinant ATF6 was further confirmed by the TG-inducible appearance of the processed ATF6 N-terminal fragment, which was blocked by the introduction of the S1/2P site mutagenesis (Fig. 1d).

**Fig. 1.**
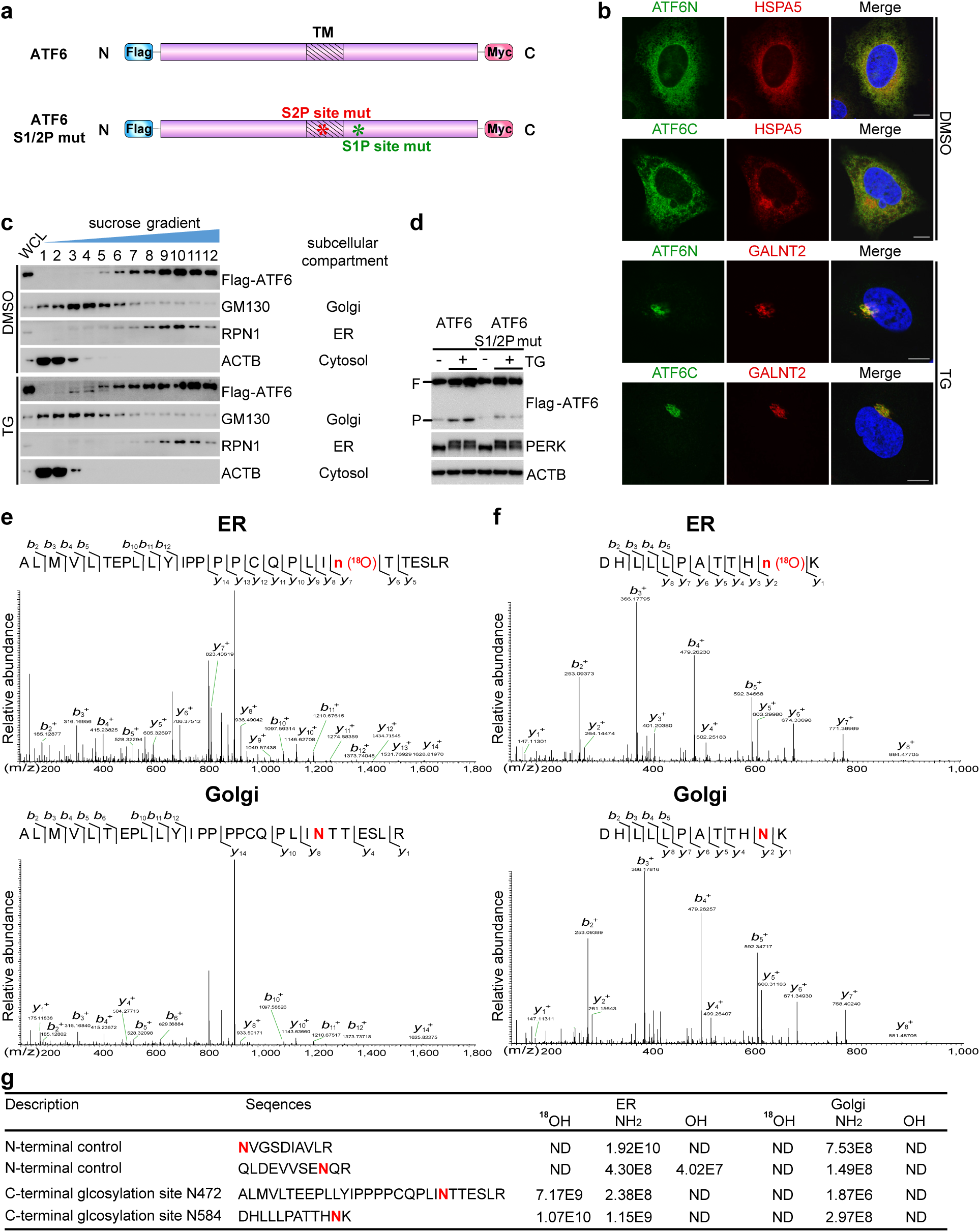
Golgi-bound ATF6 is unglycosylated rather than deglycosylated. **a**, Illustrations of wild-type and S1/2P sites-mutated ATF6 constructs. Striped boxes denote transmembrane domain. Flag and Myc tags were added to the N- and C-terminus of ATF6. **b**, IF staining of ATF6 (Flag-ATF6-Myc S1/2m). Cells were fixed with or without TG treatment (100nM, 1 h). ATF6N, Flag immunofluorescence detection of ATF6 N-terminus; ATF6C, Myc immunofluorescence detection of ATF6 C-terminus; HSPA5, heat shock protein A5, ER marker; GALNT2, Polypeptide N-acetylgalactosaminyltransferase 2, Golgi marker. Scale bar, 10 um. **c**, IB of ATF6 across discontinuous sucrose gradient fractions. Cell culture samples were collected with or without TG treatment (100nM, 5h). **d**, IB of the stress-induced (100 nM TG, 5h) processing of ATF6 (Flag-ATF6-Myc) and mutant ATF6 (Flag-ATF6-Myc S1/2m). “F”, full-length ATF6; “P”, processed ATF6. **e**, LC-MS/MS identification of ATF6 luminal domain peptide ^450^ALMVLTEEPLLYIPPPPCQPLINTTESLR^478^ from ER and Golgi fractions. The labeled peaks indicated a series of *y* or *b* ions of peptides. **f**, LC-MS/MS identification of ATF6 luminal domain peptide ^574^DHLLLPATTHNK^585^ from ER and Golgi fractions. **g**, Summary of LC-MS/MS quantifications of four Asparagine (N)-containing peptides in three different forms: Asparagine (NH_2_), deglycosylated to Aspartate in vivo (OH), deglycosylated to Aspartate in vitro (^18^OH). “ND”, not determined.

After confirming the exogenous ATF6 being able to recapitulate essential aspects of cell biology as endogenous ATF6 ^32^, we used mass spectrometry to monitor the glycosylation status of ATF6 peptides prepared from ER and Golgi fractions (Supplementary Fig. 1a). To differentiate Asparagine deglycosylation *in vitro*, and those may occur *in vivo* in response to ER stress, we introduced H_2_^18^O during PNGase F treatment of ER and Golgi samples. We predicted that ER-localized ATF6 would be glycosylated on the Asparagine residues on its C-terminus peptides, and PNGase F treatment will lead to an Asparagine to Aspartate conversion and ^18^O incorporation. If ATF6 was deglycosylated prior to stress-induced trafficking to Golgi, then Golgi-bound ATF6 should have the same Asparagine to Aspartate conversion but no ^18^O incorporation. As a control, the Asparagine residues on the N-terminus of ATF6 should remain as Asparagine as they are exposed in the cytosol and no glycosylation event should occur.

As predicted, the two Asparagine residues that have been previously reported to be glycosylated in the ER ^39^ were fully converted to ^18^O-labeled Aspartate (Fig. 1e, f, upper panels, quantified in Fig. 1g). In contrast, Asparagine residues on the ATF6 N-terminus remained as Asparagine regardless of ER and Golgi localizations (Supplementary Fig. 1b,c). Unexpectedly, none of the C-terminus Asparagine residues from the Golgi-localized ATF6 were converted to Aspartate (Figure 1e, f, bottom panels). Therefore, Golgi-bound ATF6 is newly synthesized upon stress induction, not derived from deglycosylation of pre-existing ATF6 as previously hypothesized.

### ATF6 is sorted to Golgi as a peripheral membrane protein

We applied the proteinase K protection assay to probe potential topological signatures that may drive the selective sorting of unglycosylated ATF6 to the Golgi compartment. As expected from a type II membrane protein, the ATF6 N-terminus was retained in the cytosol and readily degraded by proteinase K, regardless of TG treatment (Fig. 2a, Flag panel). However, although the ATF6 C-terminus was initially protected from proteinase K treatment (Fig. 2a, Myc panel, 0h), it became increasingly susceptible to proteinase degradation upon the onset of TG treatment (Fig. 2a, Myc panel, 2-8h, quantified in Fig. 2b), suggesting externalization of its C-terminus in the cytosol. As a control, the ER-resident protein PERK and the Golgi-resident MAN2A1 remained protected throughout TG treatment (Fig. 2a, PERK and MAN2A1 panels). We further separated ER and Golgi-bound ATF6 through sucrose gradient fractionation and probed their respective topology. As shown in Fig. 2c, the C-terminus of ER-bound ATF6 was protected from proteinase degradation in the absence of membrane-permeating reagent TX-100, whereas the Golgi-bound ATF6 was readily degraded.

**Fig. 2.**
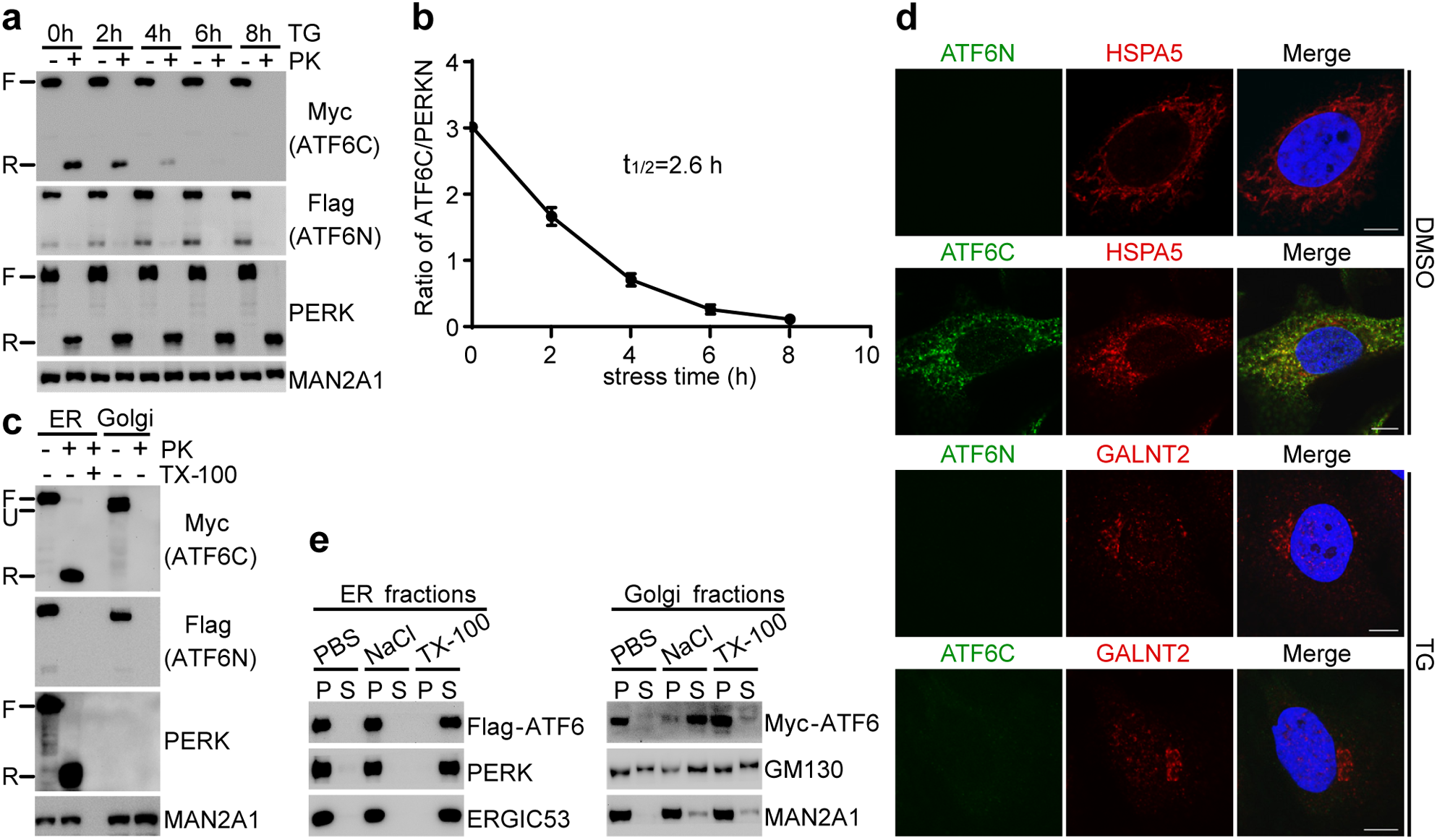
ATF6 is peripherally associated with the Golgi under ER stress conditions. **a**, Proteinase K protection assay of membrane fractions prepared from control and stressed (20nM TG) cell samples. “F”, full-length of indicated proteins; “R”, remnant signals of the luminal fragment of corresponding proteins protected from proteinase K degradation. **b**, Quantification of ATF6 C-terminus (ATF6C) remnant signals measured by Myc immunoblot, normalized to the remnant signals of PERK N-terminus (PERKN) from (**a**). t_1/2_ means the half life of normalized ATF6C remnant signal. Data are presented as means ± SEM, n=3. **c**, Proteinase K protection assay of ER and Golgi fractions prepared from stressed (100nM TG, 5h) cell samples. “U”, unglycosylated ATF6. **d**, Fluorescence proteinase protection (FPP) assay of normal and stressed (100nM TG, 1 h) cells. Scale bar, 10 um. **e**, Topology analysis of ER- and Golgi-bound ATF6 with salt wash (0.5 M NaCl) and detergent treatment (2% TX-100). PERK, ERGIC53, and MAN2A1, markers for integral membrane protein; GM130, peripheral membrane protein marker.

To avoid potential artifacts that may be introduced during the process of membrane isolation, we examined ATF6 topology *in situ* with fluorescence proteinase K protection assay (Fig. 2d). Consistent with the membrane fractionation assay, the ATF6 C-terminus, but not its N-terminus, was protected from proteinase treatment under normal, non-stress conditions (Fig. 2d, DMSO panel, and compare with Fig. 1b, DMSO panel). In contrast, the fluorescent signals marking both ends of the Golgi-bound ATF6 were lost after proteinase treatment (Fig. 2d, TG panel, compare with pre-proteinase treatment Fig. 1b, TG panel), confirming the exposure of Golgi-bound ATF6 C-terminus in the cytosol.

As ATF6 harbors a single transmembrane at the center of its peptide sequence, the stress-induced retention of both of its N- and C-terminus in the cytosol suggests it may be peripherally associated with the Golgi membrane. Indeed, the Golgi-bound ATF6 was readily stripped by salt wash as another peripheral protein, GM130 (Fig. 2e, Golgi fractions). Intriguingly, the Golgi-bound ATF6 was resistant to TX-100 treatment to a similar degree as another Golgi peripheral membrane protein GM130 and the integral membrane protein MAN2A1; both were reported to be detergent-resistant ^41,42^. In contrast, the ER-bound ATF6 could only be released by the membrane dissolving detergent TX-100 (Fig. 2e, ER fractions), consistent with it being a type II integral membrane protein on the ER.

### Genetic and chemical inhibition of the SEC61 translocon promotes ATF6 sorting to the Golgi

The stress-induced peripheralization of ATF6 explains why its Golgi-bound form is unglycosylated. However, is ATF6 peripheralization sufficient to drive its Golgi trafficking? We sought to identify and manipulate biological components involved in ATF6 biogenesis to examine the role of impaired membrane integration in regulating its ER-to-Golgi trafficking. There are two types of protein machineries in membrane protein biogenesis: the SEC61 translocon and the WRB /CAML ^43,44^. Reducing the SEC61 translocon levels by small interference RNA (*siSEC61A1*) potently suppressed the biogenesis of ATF6, and almost half of them appeared in a fast-migrating and presumably non-glycosylated form even under non-stress conditions (Fig. 3a). Crude fractionation of cell lysates showed that ATF6 in the heavy membrane fraction (M) was mostly glycosylated, whereas the light membrane and cytosolic fraction (C) was exclusively represented by fast-migrating, non-glycosylated ATF6 (Supplementary Fig. 2a). Refined fractionation through sucrose gradient revealed that the vast majority of the fast-migrating ATF6 was not in the cytosol but the Golgi fractions (Fig. 3b and Supplementary Fig. 2b), and the Golgi-bound ATF6 induced by *SEC61A1* knockdown had the same peripheral topology as TG treatment (Fig. 3c). Immunofluorescence studies further confirmed the concentration of ATF6 in the Golgi compartment and the cytosolic exposure of its N- and C-terminus in the SEC61 knockdown cells (Fig. 3d,e). The effect of WRB complex knockdown on ATF6 distribution was also examined, and its impact on ATF6 Golgi-targeting was much less pronounced (Fig. 3a,b,d,e, Supplementary Fig. 2a,b).

**Fig. 3.**
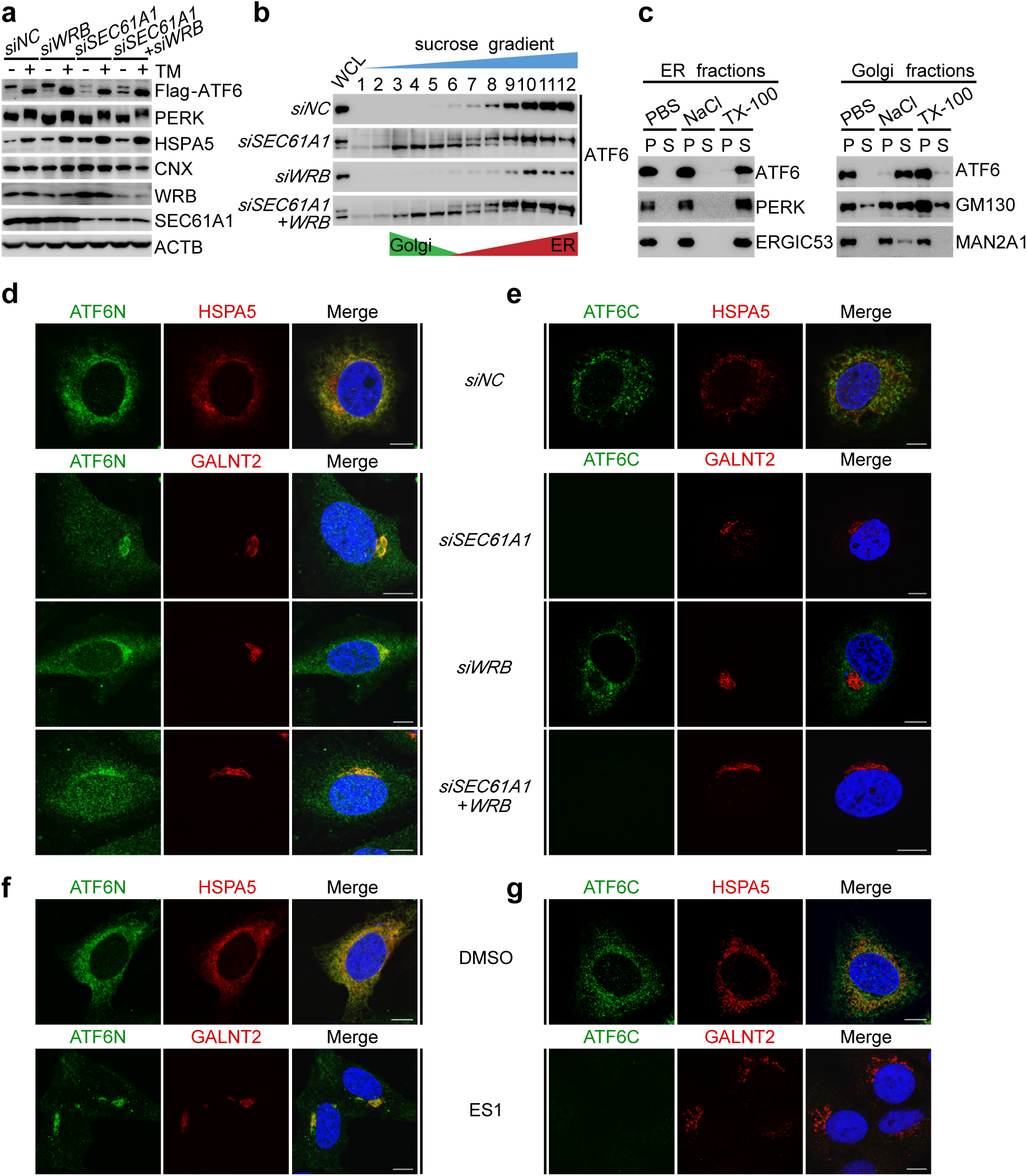
Translocon blockage promotes ATF6 translocation to the Golgi. **a**, IB of ATF6 and other ER resident proteins in control and target gene-knockdown cells with (5ug/ml Tm, 5h) or without stress induction. **b**, IB of ATF6 (Flag) across discontinuous sucrose gradient fractions prepared from control and target gene-silenced cells without stress induction. **c**, Topology analysis of ER- and Golgi-bound ATF6 from (**b**) with salt wash (0.5 M NaCl) and detergent treatment (2% TX-100). ATF6 was detected by Flag Ab in ER fractions and by Myc Ab in Golgi fractions. **d**, IF of ATF6 (Flag-ATF6-Myc S1/2m). Cells were transfected with scramble or target gene siRNA for 24hs before fixation. Scale bar, 10 um. **e**, FPP assay of ATF6 C-terminus (Myc). Cells were transfected with scramble or target gene siRNA for 24hs before being fixed. Scale bar, 10 um. **f**, IF of ATF6 (Flag-ATF6-Myc S1/2m) in control (DMSO) and ES1-treated (20nM, 2h) cells. Scale bar, 10 um. **g**, FPP assay of ATF6 detected by its C-terminus Myc epitope. Cells were fixed with or without ES1 treatment (20nM, 2h). Scale bar, 10 um.

To avoid long-term and potentially indirect effects caused by SEC61 knockdown, we asked whether acute, chemical inhibition of the translocon complex may have the same effect. Indeed, acute SEC61 translocon blockage by the chemical inhibitor Eeyerastatin 1 (ES1) led to rapid partitioning of ATF6 to the Golgi compartment within 5 hours (Fig. 3f and Supplementary Fig. 2c), which was accompanied by the appearance of the processed form (Supplementary Fig. 2d), and the induction of ATF6 target genes (Supplementary Fig. 2e). The Golgi-bound ATF6 under ES1 treatment conditions adopted the same peripheral membrane protein topology as evidenced by its susceptibility to proteinase K treatment (Fig. 3g and Supplementary Fig. 2f) and salt extraction (Supplementary Fig. 2g). Therefore, reducing cellular translocon capacity by both chemical and genetic means seems to be sufficient to drive ER-to-Golgi trafficking and activation of ATF6.

### ATF6 transmembrane domain and C-terminus regulate membrane insertion and trafficking

ATF6 is a type II transmembrane protein with a centrally-placed transmembrane domain (Supplementary Fig. 3a), and its proper insertion into the ER membrane requires the full translocation of its C-terminus (∼270aa) into the ER lumen. We constructed a series of mutants covering the transmembrane domain and its C-terminus to dissect intrinsic factors that may enable it to sense ER stress and modulate its membrane insertion efficiency.

We first confirmed the importance of the transmembrane by replacing a stretch of hydrophobic residues (^387^FILL^390^) with positively charged Arginine residues (387-390R, Supplementary Fig. 3a). Disruption of this α-helical transmembrane domain led to a loss of ER localization (Supplementary Fig. 3b) and ER stress-induced activation (Supplementary Fig. 3c). Replacing the single-pass transmembrane domain with the dual-pass hairpin structure from DGAT2 enabled the ATF6 chimera protein to be properly localized to the ER as an integral membrane protein with both of N- and C-terminus exposed in the cytosol (Supplementary Fig. 3d-f). However, TG treatment did not impair the insertion of the ATF6(DGAT2) chimera protein into the ER membrane, as evident for its resistance to salt extraction (Supplementary Fig. 3g, TG panel), and not partitioned to the Golgi compartment (Supplementary Fig. 3e,h). Together, these results suggest the ATF6 transmembrane domain regulates ER-targeting, whereas the cross-membrane translocation of its C-terminus serves as a regulatory node in stress sensing.

Next, we performed domain swapping experiments between ATF6 and its ER stress insensitive homologue Luman to examine the role of ATF6 C-terminus in regulating membrane insertion and Golgi trafficking (Fig. 4a). As demonstrated in Fig. 4b, replacing the N-terminus of ATF6 with its counterpart from Luman did not affect the ability of the chimera protein (Luman-ATF6) to sense ER stress by translocating to the Golgi compartment. In contrast, substituting the ATF6 C-terminus with its much shorter Luman counterpart (ATF6-Luman) completely abrogated its ability to sense ER stress (Fig. 4b, quantified in Fig. 4c). More importantly, the ability of the chimera proteins to sense ER stress was strictly and negatively correlated with its membrane insertion efficiency: TG treatment led to a rapid exposure of the C-terminus of the Luman-ATF6 chimera protein but not the ATF6-Luman chimera (Fig. 4d,e, and quantified in 4f).

**Fig. 4.**
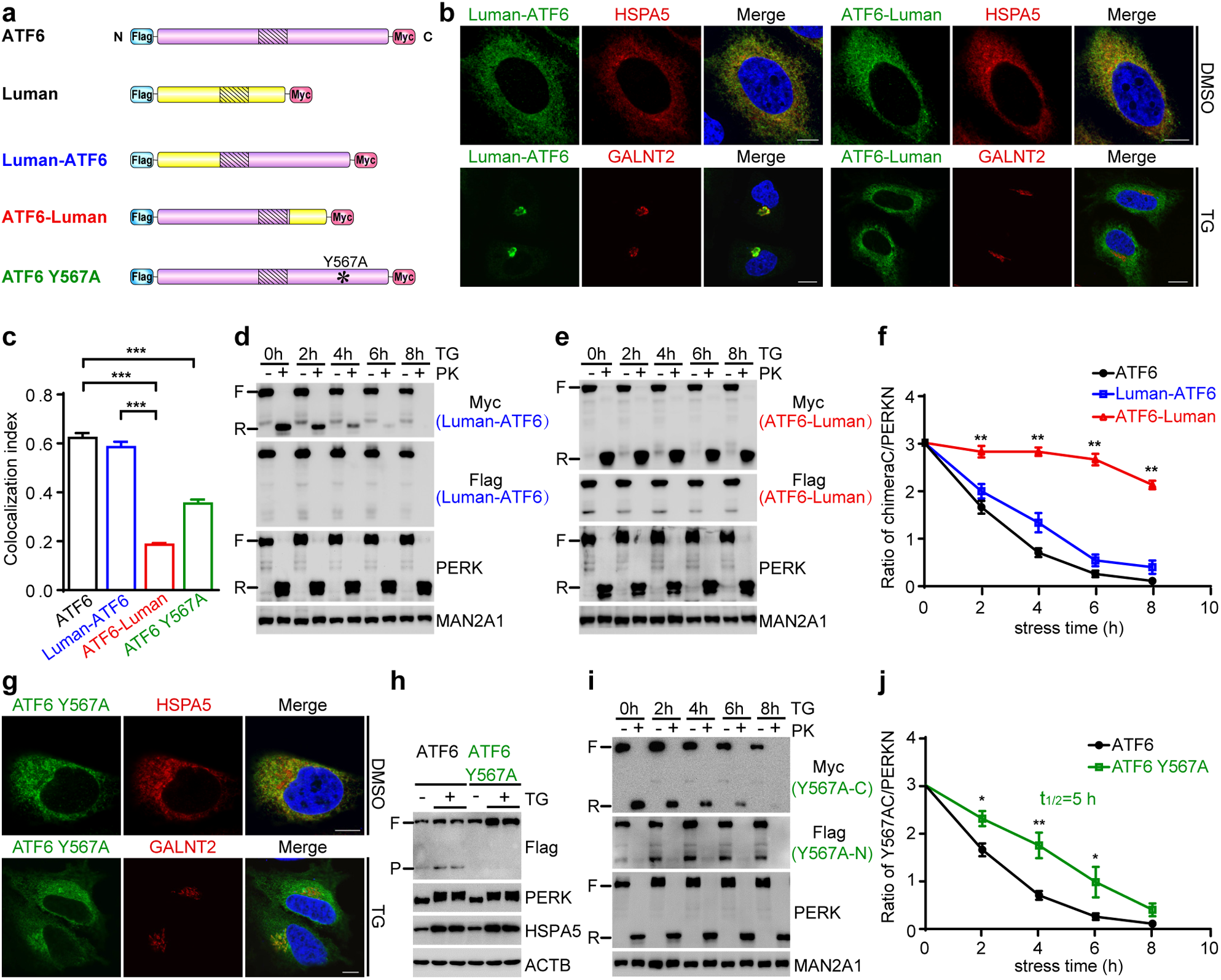
ATF6 membrane insertion efficiency is controlled by its transmembrane domain and C-terminus. **a**, Illustrations of ATF6 (purple), Luman (yellow), chimera proteins (Luman-ATF6, ATF6-Luman) and Y567A mutant of ATF6. **b**, IF staining of chimera proteins (Flag). Cells were transfected with scramble siRNA for 24hs and fixed with or without prior ER stress induction (100nM TG, 1h), related to **Supplementary Fig. 5b**. Scale bar, 10 um. **c**, Pearson’s correlation coefficiency of indicated proteins with GALNT2 under TG treatment (100nM, 1h) conditions. Data are means ± SEM, n=100, ***p < 0.001, Student’s t-test. **d**,**e**, Proteinase K protection assay of chimera proteins (**d**: Luman-ATF6, **e**: ATF6-Luman). Cell culture samples were collected before and during the course of 8-hr 20nM TG treatment. **f**, Quantification of remnant of Myc signals normalized to PERKN from (**d**) and (**e**), and compared with wild-type ATF6 in **Fig. 2b**. Data are means ± SEM, n=3, *p < 0.05; **p < 0.01, two-way ANOVA. **g**, IF staining of ATF6 Y567A (Flag). Cells were transfected with scramble siRNA for 24hs and fixed with or without prior ER stress induction (100nM TG, 1h), related to **Supplementary Fig. 5c**. Scale bar, 10 um. **h**, IB of the stress-induced (100 nM TG, 5h) processing of ATF6 and ATF6 Y567A. **i**, Proteinase K protection assay of ATF6 Y567A. Cell culture samples were collected prior to and during TG treatment (20nM). **j**, Quantification of remnant ATF6 Y567A (Myc) normalized to PERKN in (**i**), and compared with wild type ATF6 in **Fig. 2b**. Data are means ± SEM, n=3, p < 0.05; **p < 0.01, two-way ANOVA.

Recent reports suggested a single point mutation (Y567N) derived from human achromatopsia patients was able to impair ATF6 trafficking to the Golgi ^45^. We suspected that the bulk size of the tyrosine residue in the context of a stretch of charged residues (^560^RRRGDTFYVVSFRRDH^575^) might pose a challenge for translocon import under stress conditions. To test this hypothesis, we mutated Y567 to Alanine (Y567A) and found that it strongly suppressed the stress-induced ER-to-Golgi trafficking (Fig. 4g, quantified in 4c) and processing (Fig. 4h). Importantly, the inability of the mutant protein to respond to ER stress was correlated with improved cross-membrane translocation of the ATF6 C-terminus and increased resistance to proteinase K degradation during TG treatment (Fig. 4i, quantified in Fig. 4j). Therefore, not only the length but also sequence signatures present in the ATF6 C-terminus may regulate its membrane insertion efficiency and stress responsiveness.

### The BAG6 complex recognizes ATF6 for membrane targeting and trafficking

The inability of the ATF6(DGAT2) chimera protein to be partitioned to Golgi suggests that the exposure of ATF6 C-terminus is not sufficient for it to be recognized and delivered to the COPII vesicles. Therefore, we sought to identify protein machinery that may recognize and regulate ATF6 partitioning.

As ATF6 integration into the ER membrane depends primarily on the SEC61 translocon, we first evaluated the involvement of SRP54 (signal recognition particle 54 kDa protein), an established upstream component of the translocon pathway, in ATF6 membrane targeting. Unexpectedly, knocking down the expression of SRP54 did not affect ATF6 localization to the ER and Golgi under both normal and stress conditions (Supplementary Fig. 4a,b), suggesting the involvement of novel receptor proteins in recognizing and delivering ATF6 to the membrane system. Therefore, we performed co-immunoprecipitation of ATF6 recombinant proteins followed by mass spectrometry to identify such candidate proteins (Table S1). We focused on BAG6 (BCL2-associated athanogene 6) because its interaction with ATF6 was significantly increased under ER stress conditions (Fig. 5a), and it has the demonstrated capacity to triage client proteins between membrane insertion and degradation pathways ^46^.

**Fig. 5.**
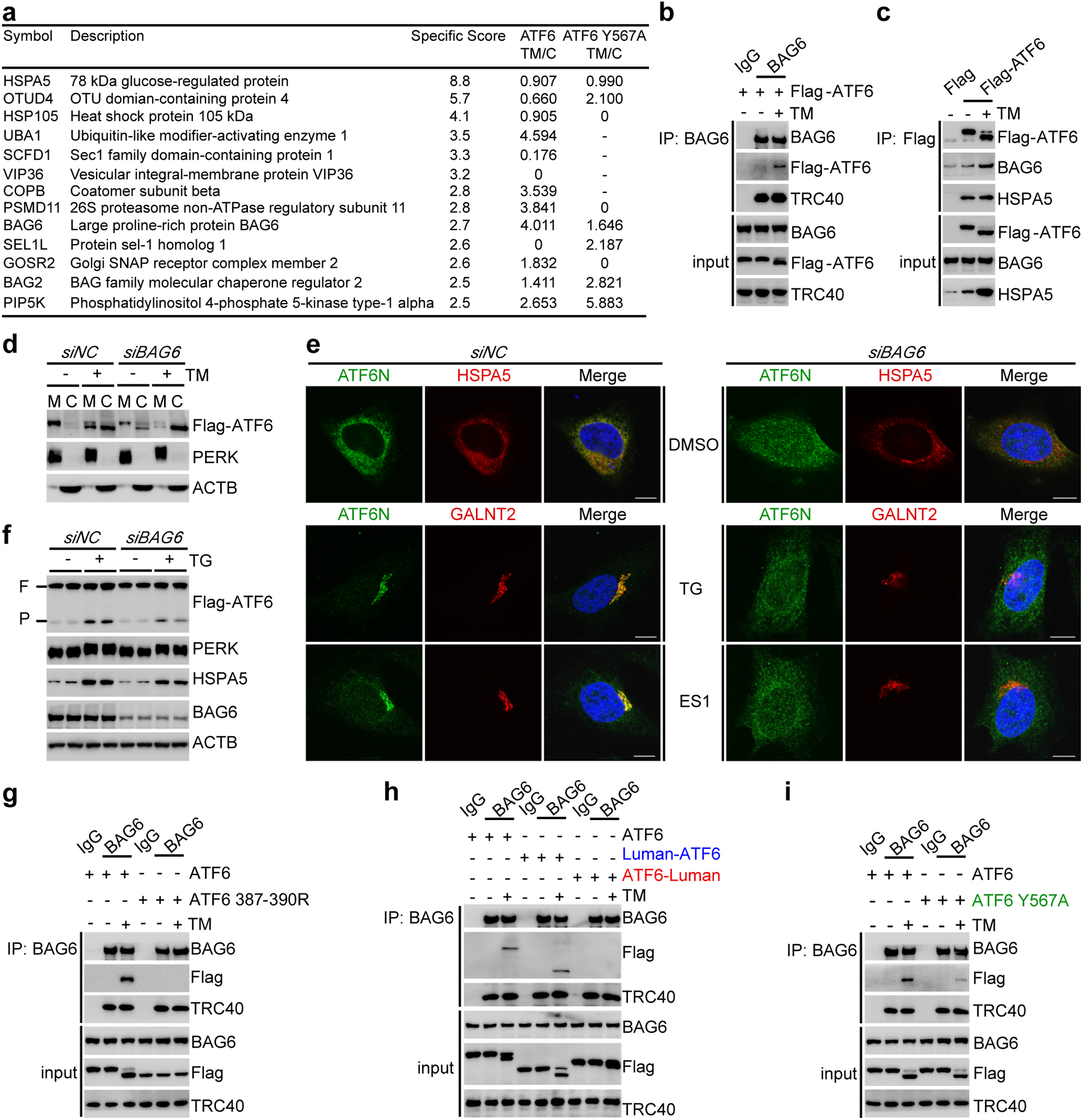
The BAG6 complex is required for ATF6 membrane insertion and Golgi targeting. **a**, List of ATF6 binding proteins in MS. “Specificity Score”, fold-of-enrichment in ATF6 pulldown over the negative control; “TM/C”, enrichment of ATF6 pulldown in TM treated samples over DMSO controls (C); “0”, score missing in TM samples; “-”, score missing in control samples. **b**, Co-IP of ATF6 by BAG6 antibody under control and stress (5 ug/ml TM, 5h) conditions. **c**, Co-IP of BAG6 by ATF6 (Flag beads) under control and stress (5 ug/ml TM, 5h) conditions. **d**, IB of ATF6 in heavy membrane (M) and cytosol fractions (C, include light membrane). Cells were transfected with either scramble or *BAG6* siRNA for 24hs and collected with or without prior TM treatment (5ug/ml, 5h). **e**, IF staining of ATF6 (Flag). Cells were transfected with either scramble or *BAG6* siRNA for 24hs and fixed with or without indicated chemical treatment (100nM TG, 1h; 20nM ES1, 2h). Scale bar, 10 um. **f**, IB of ATF6 processing. Cells were transfected with either scramble or *BAG6* siRNA for 24hs and collected with or without prior TG treatment (100nM, 5h). **g**, Co-IP of ATF6 387-390R by BAG6 antibody. Cell culture samples were collected after 5hs of DMSO or TM (5ug/ml) treatment. **h**, Co-IP of ATF6 chimera proteins by BAG6. **i**, Co-IP of ATF6 Y567A by BAG6.

We were able to confirm the interaction between ATF6 and BAG6 under both normal and stress conditions by co-immunoprecipitation (Fig. 5b,c), and the stabilization of ATF6-BAG6 interaction was correlated with the accumulation of the non-glycosylated form of ATF6 and its trafficking to the Golgi compartment (Fig. 5b,c and 1c). Silencing the expression of BAG6 complex reduced the protein level of ATF6 (Supplementary Fig. 5a) and resulted in the appearance of the non-glycosylated form of ATF6 in the cytosol (Fig. 5d). ATF6 became dispersed throughout the BAG6 complex knockdown cells (Fig. 5e, right panel). Further treatment of cells with TG or ES1 failed to induce apparent Golgi translocation, and the level of ATF6 processing was significantly reduced (Fig. 5e,f). Together, these results identified BAG6 complex as a novel receptor system upstream of the SEC61 translocon and the ER-to-Golgi vesicle trafficking system in regulating ATF6 biogenesis.

The BAG6 complex primarily function is as a receptor for tail-anchored (TA) membrane proteins, and extended retention of client proteins with BAG6 usually leads to their degradation ^46,47^. However, ATF6 represents a novel substrate for BAG6 because it harbors a centrally-located transmembrane domain. Substituting the hydrophobic residues in the transmembrane domain with positively-charged Arginine residues (ATF6 387-390R) disrupted BAG6 binding, ER localization, and stress-induced activation (Fig. 5g and Supplementary Fig. 3b,c). In contrast, mutations in the N- and C-terminus of ATF6 did not affect their BAG6-dependent ER localization (Supplementary Fig. 5b,c). Strikingly, mutations that modulated the extent of ATF6 retention with BAG6 were strictly correlated with their Golgi partitioning efficiency. As shown in Fig. 5h, the Luman-ATF6 chimera protein that was readily sorted to Golgi exhibited extended interaction with BAG6 under ER stress conditions, whereas the ATF6-Luman protein with no ER stress response capability showed no stress-induced retention with the BAG6 complex. The single point mutant (ATF6 Y567A) with reduced capacity to respond to ER stress also showed less retention with BAG6 (Fig. 5i). Therefore, our results suggest a novel function of BAG6 system in triaging ATF6 between membrane insertion and secretory pathways that is dependent upon translocon efficiency and stress levels.

### The protease domain of the Golgi-bound S1P resides in the cytosol

Our finding that Golgi-bound ATF6 is a peripheral membrane protein would require its protease S1P to cleave ATF6 on the cytosolic side of Golgi, which is incompatible with the established type I topology of S1P on the ER ^32,48,49^. Therefore, we speculated that S1P might adopt a new topology under ER stress conditions due to the potential impairment in the cross-membrane translocation of its N-terminus (∼1000aa).

To evaluate this hypothesis, we constructed a cell line that stably expressed a low level of exogenous S1P protein, and we inserted three epitope tags to monitor its topology under normal and stress conditions (Fig. 6a). Under normal conditions, S1P was highly glycosylated and migrated between 130-180kDa (Supplementary Fig. 6). Treatment of cells with TG led to the appearance of a faster-migrating S1P with an apparent molecular weight of less than 130kDa (Supplementary Fig. 6), possibly an under- or un-glycosylated form of S1P. Sucrose gradient fraction showed that the faster-migrating form of S1P was specifically co-fractionated with the Golgi marker GM130 (Fig. 6b).

**Fig. 6.**
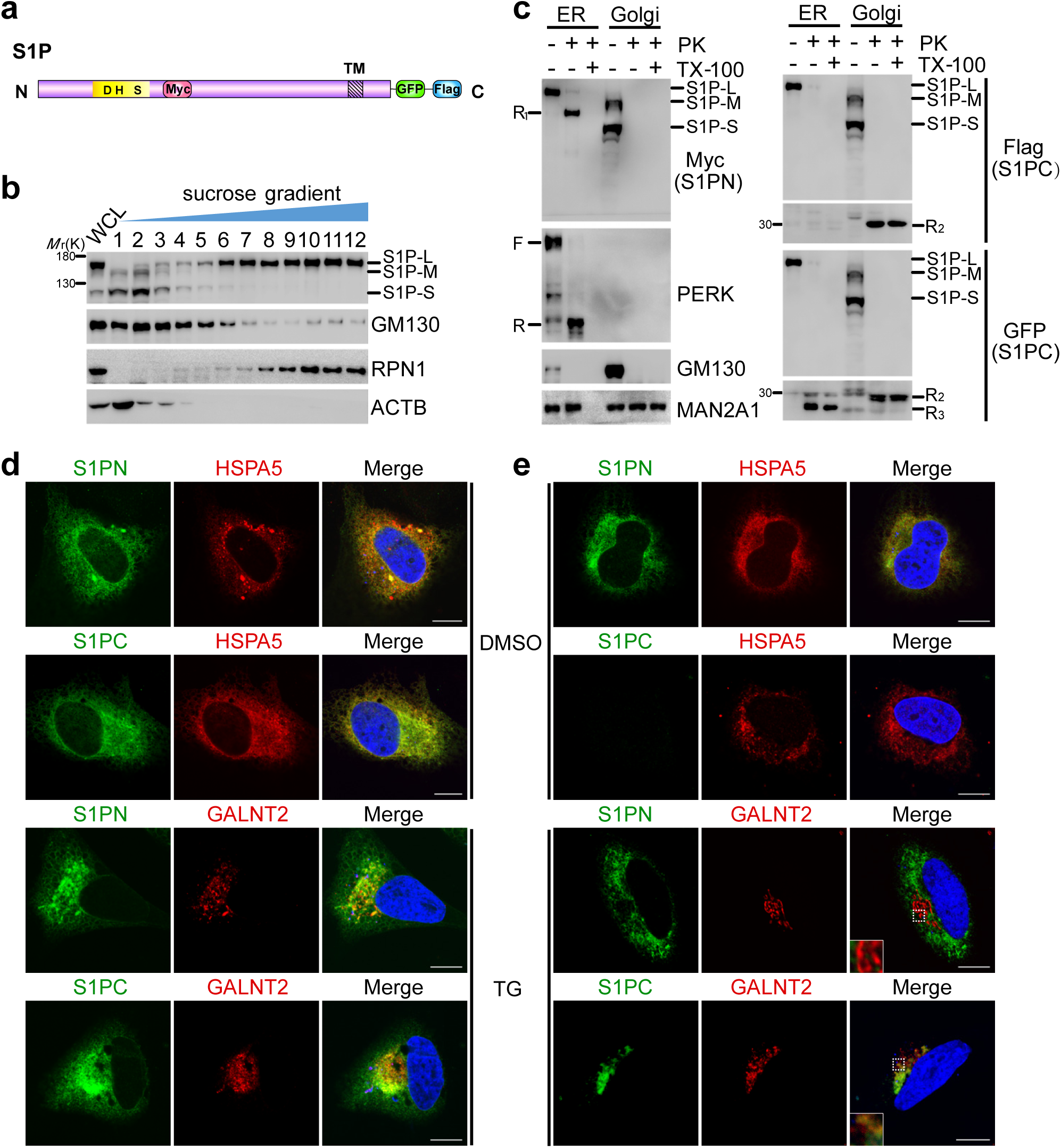
Distinct topologies of S1P on the ER and Golgi. **a**, Illustration of S1P constructs. Three epitope tags were incorporated to monitor S1P topology. The protease domain was marked by its catalytic residues D, H, and S. **b**, IB of S1P (Flag) across sucrose gradient fractions. Cell culture samples expressing low levels of the exogenous S1P were collected after 10hs of TG treatment (100nM). “L”, “M”, “S”, S1P of different sizes. **c**, Proteinase K protection assay of S1P. ER and Golgi fractions were prepared from (**b**). “R”, remnant fragment of PERK (PERKN); “R_1_”, remnant fragment of S1P with C-terminus degraded; “R_2_”, remnant fragment of S1P with N-terminus degraded; “R_3_”, residual GFP fragment. **d**, IF staining of S1P (Myc-S1P-Flag) in control and stressed (100nM TG, 10 h) cells. Scale bar, 10 um. **e**, FPP assay of S1P in control and stressed (100nM TG, 10 h) cells. Scale bar, 10 um.

As expected, the catalytic domain of ER-bound S1P as detected by the Myc tag was resided in the ER lumen and protected from proteinase K (Fig. 6c, left panel, “ER” lanes). In contrast, the catalytic domain of the Golgi-bound S1P was readily degraded by proteinase K treatment even in the absence of membrane permeabilizing agent TX-100 (Fig. 6c, left panel, “Golgi” lanes). The topological difference between ER and Golgi-bound S1P was not limited to its N-terminus. As shown on the right panel of Fig. 6c, proteinase K treatment fully degraded the Flag epitope signal marking the C-terminus of ER-bound S1P (Fig. 6c, right/top panel, “ER” lanes) with the GFP module being resistant to proteinase degradation (Fig. 6c, right/bottom panel, “ER” lanes, R3 fragment), but a ∼30kDa fragment remained for its Golgi-bound counterpart (Fig. 6c, right panel, “Golgi” lanes, R2 fragment) that contained both the GFP and the Flag epitopes. Therefore, the much shorter C-terminus of S1P, instead of its bulky N-terminus, is transported into the ER lumen and sorted to Golgi under stress conditions.

The stress-induced type I to type II topology change of S1P was further confirmed by immunofluorescence. As shown in Fig. 6d, S1P primarily localized to the ER compartment under normal conditions (Fig. 6d, DMSO panel), and the immunofluorescent signal of the C-terminal (S1PC), but not N-terminal (S1PN), was readily degraded by proteinase treatment (Fig. 6e, DMSO panel), confirming its type I topology. TG treatment resulted in enrichment of S1P in the Golgi compartment labeled by GALNT2 immunofluorescence (Fig. 6d, TG panel), suggesting a stress-enhanced sorting of S1P to the Golgi in a similar manner to ATF6. More importantly, ER stress caused specific loss of S1PN immunofluorescence signal in the Golgi but not in the ER compartment after proteinase treatment, whereas the S1PC signal on the Golgi compartment was fully resistant to proteinase degradation (Fig. 6e, TG panel).

Therefore, stress-induced interference of membrane protein insertion is not restricted to ATF6; correspondingly changes in the S1P protease topology ensure ATF6 can be properly processed and activated on the cytosolic side of the Golgi compartment.

### Translocon inhibition expedites ER-to-Golgi trafficking and activation of SREBP2

S1P processes many substrates in addition to ATF6 on the Golgi. The topology-driven sorting of ATF6 and S1P observed above led us to ask whether such a mechanism may be extrapolated to the regulated trafficking of other ER-tethered transcription factors. SREBP2 is a canonical ER-bound transcription factor that is activated by S1P through regulated trafficking in response to cholesterol depletion ^50^. The exogenously expressed SREBP2 was able to respond to cholesterol depletion caused by serum starvation by enhanced production of the N-terminus, transcriptionally active nucleus fragment (Fig. 7a), which was blocked by mutating the S1/2P cleavage site (Supplementary Fig. 7a). Similar to the effect of ER stress on interfering ATF6 membrane insertion, cholesterol depletion reduced membrane integration of the SREBP2 protein, as demonstrated by its increased retention in the cytosol upon serum starvation (Fig. 7b). The membrane biogenesis of SREBP2 also required the SEC61 translocon, but not the BAG6 chaperone complex (Supplementary Fig. 7b), as SEC61 knockdown reduced SREBP2 incorporation into the membrane fraction (Fig. 7b). Chemical (ES1) or genetic (*siSEC61A1*) inhibition of translocon function effectively accelerated SREBP2 sorting and activation under serum starvation conditions (Fig. 7c-e), although blocking translocon function alone was not sufficient (Supplementary Fig. 7c,d). Therefore, the regulated ER-to-Golgi trafficking and activation of SREBP2 are also accompanied by altered membrane integration.

**Fig. 7.**
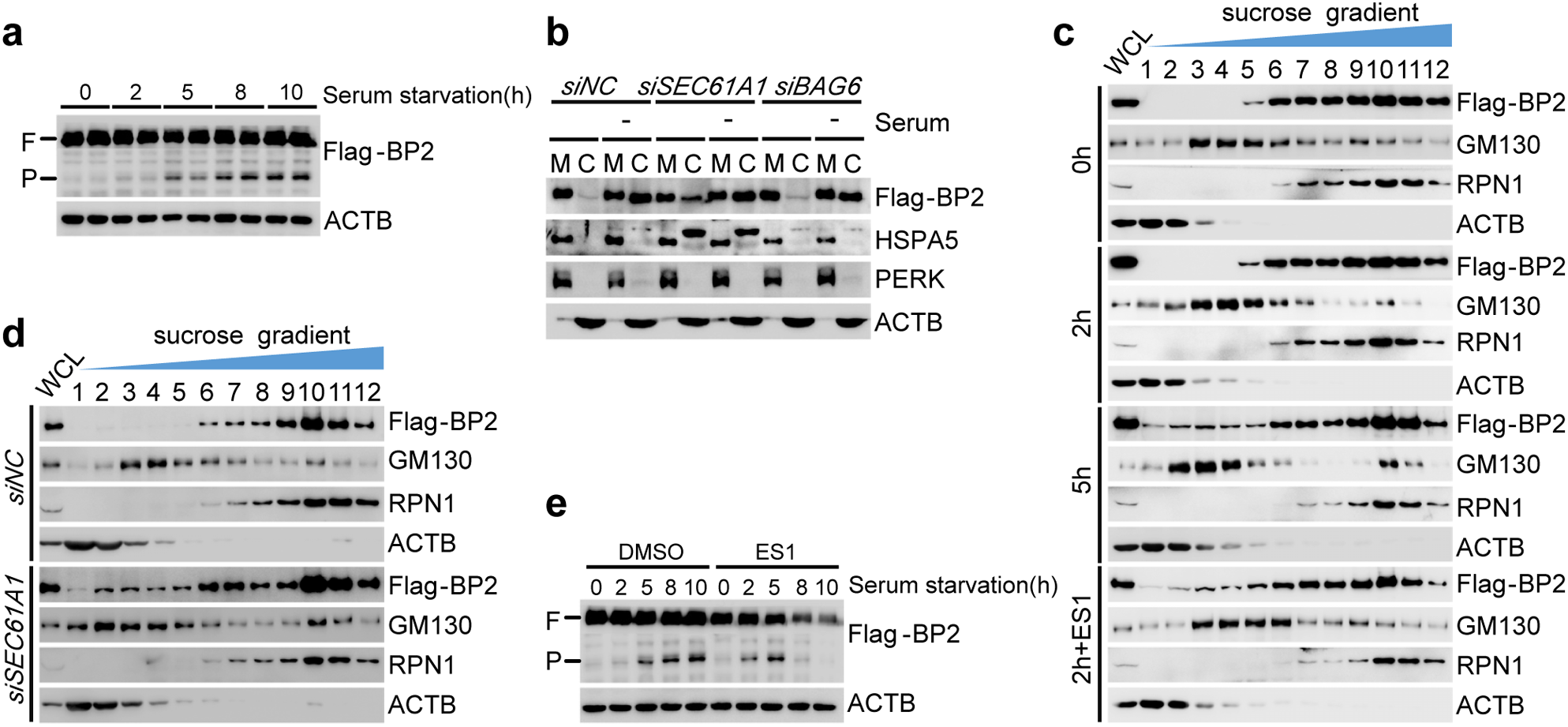
Translocon inhibition accelerates SREBP2 activation during serum starvation. **a**, IB of SREBP2. Stable cell line expressing low levels of Flag-tagged SREBP2 were subjected to serum withdraw, and cell culture samples were collected at indicated time points. **b**, IB of SREBP2 from heavy membrane (M) and cytosol (C, include light membranes) fractions. Cells were transfected with siRNA targeting indicated genes for 24hs before subjected to 5hs of serum withdrawal. “-”, serum starvation. **c**, IB of SREBP2 across sucrose gradient fractions. Cell culture samples were collected after 0, 2, 5hs of serum withdrawal, or 2hs of serum withdrawal plus 20nM of ES1 co-treatment. **d**, IB of SREBP2 across sucrose gradient fractions. Cells were transfected with indicated siRNAs for 24hs before subjecting to 2hs of serum withdrawal. **e**, IB of SREBP2 processing. ES1 co-treatment at 20nM.

## Discussion

Here we show that stress-induced activation of ATF6 is driven by interfering its membrane integration process rather than the release of pre-existing proteins upon the initiation of ER stress. Our findings are consistent with previous observations that Golgi-bound ATF6 is under-glycosylated and reduced, although those findings were misinterpreted as the result of deglycosylation and reduction due to the lack of topological information ^39,51^. Presently, there is insufficient information to conclude the accelerated Golgi sorting and activation of SREBP2 by translocon inhibition is due to conformational changes in SREBP2. However, cholesterol-induced conformational change of SCAP has been well demonstrated ^52–55^. Additionally, the cytosolic location of the Golgi-bound S1P catalytic domain strongly suggests the cleavage sites of its substrate proteins are similarly localized in the cytosol, and thus such a topology-driven sorting and activation mechanism is broadly employed in the cell. Therefore, we propose a sensing-by-synthesis model, in which cellular sensors are constantly synthesized, captured, and degraded under homeostatic conditions, and perturbations in the cellular and ER environment that may prevent the proper ER membrane integration of a specific sensor will cause it to be sorted to the Golgi and activated.

Compared to the release-from-retainer model, sensing-by-synthesis is more broad and robust, by not relying on a single retainer protein. In the case of ATF6, the revised model allows broad surveillance of all biochemical and cellular processes involved in ATF6 biogenesis that may be perturbed by ER stress, not just BiP/GRP78 abundance. The current model of ATF6 activation-by-BiP-dissociation would also require the limited number of ATF6 protein to monitor the status of chaperones and misfolded proteins that are thousands of fold in excess. In contrast, sensing-by-synthesis enables ATF6 to monitor translocon function at a much lower stoichiometric ratio, because the number of translocons in each cell is much more limited ^56–58^. Although the membrane topology of the other two UPR sensors, PERK and IRE1, is not sensitive to ER stress (Figure S2A), possibly due to their slow turnover rates, the observation that both of them are localized in proximity to the translocon ^59,60^ may allow them to survey translocon function with a similar capacity.

A critical constraint for the sensing-by-synthesis mechanism is the turnover rate of the sensor itself. Work from the Lee’s and the Mori’s groups showed that ATF6 is rapidly synthesized and degraded ^37,39,40^, and therefore is compatible with such a sensing-by-synthesis mechanism. However, as all of our observations were made under conditions that ATF6 was activated in a manner of hours, the much more rapid induction of ATF6 by DTT treatment could be activated through the classical model ^34^. Future studies are required to evaluate the contribution of these different mechanisms under physiological and pathological conditions.

As newly synthesized and pre-existing ATF6 primarily differ in their glycosylation status (unglycosylated *vs.* deglycosylated), extrapolation of our model to other ER-tethered transcription factors will require knowledge about their glycosylation status in the ER and Golgi. This may be more challenging for polytopic membrane proteins like SREBP2 and SCAP, as the exact topological perturbation caused by cholesterol depletion remains to be completely defined ^53,61^. Also, the molecular mechanism allowing the topologically-altered ATF6 to be recognized and packaged into the COPII vesicles is unknown. There is an intriguing possibility that the COPII machinery may be utilized by the BAG6 system to dispose misfolded proteins with hydrophobic motifs ^62–64^, and this mechanism is adopted by the UPR for ATF6 activation. Candidate proteins identified from mass spectrometry (Figure 5A) may provide an entry point toward elucidating the biochemical mechanisms linking BAG6 to the COPII machinery. Alternatively, there may be sequence motifs in the ATF6 C-terminus that enables its recognition by COPII receptors in a manner similar to that of SREBP-SCAP ^52,55^. A large number of ATF6 mutants, either through experimental mutagenesis, or derived from human patients, have been characterized for their ability to regulate ER-to-Golgi trafficking ^28,35,40,65^. These mutant constructs may be used to capture interacting proteins that regulate ATF6 trafficking and serve as a springboard in elucidating the general mechanism governing the activation of ER-tethered transcription factors.

## Acknowledgments

We thank Drs. Yeguang Chen and Li Yu for careful reading of the manuscript. We thank the technical assistance of Yanli Zhang and Jinyu Wang (Cell Imaging Facility) at the Center for Biomedical Analysis and the Technology Center for Protein Research, Tsinghua University. We thank Gong Zhang and Peng R. Chen (Peking University) for technical suggestions and reagent sharing. We also thank Fangfang Sun for guidance in data analysis and all other members of the Fu lab for insightful discussions. The work is supported by National Science and Technology Major Project (2016YFA0502002, 2017YFA0504603), National Natural Science Foundation of China (NSFC 81471072 and 31671229), National 1000 Junior Scholar Program, and the Tsinghua-Peking Center for Life Sciences.

## Author information

Conceptualization: J.X and S.F; Investigation: J.X, X.M, F.W; Writing: J.X, S.F; Supervision: S.F, H.D.

---

**Supplementary Fig. 1.**
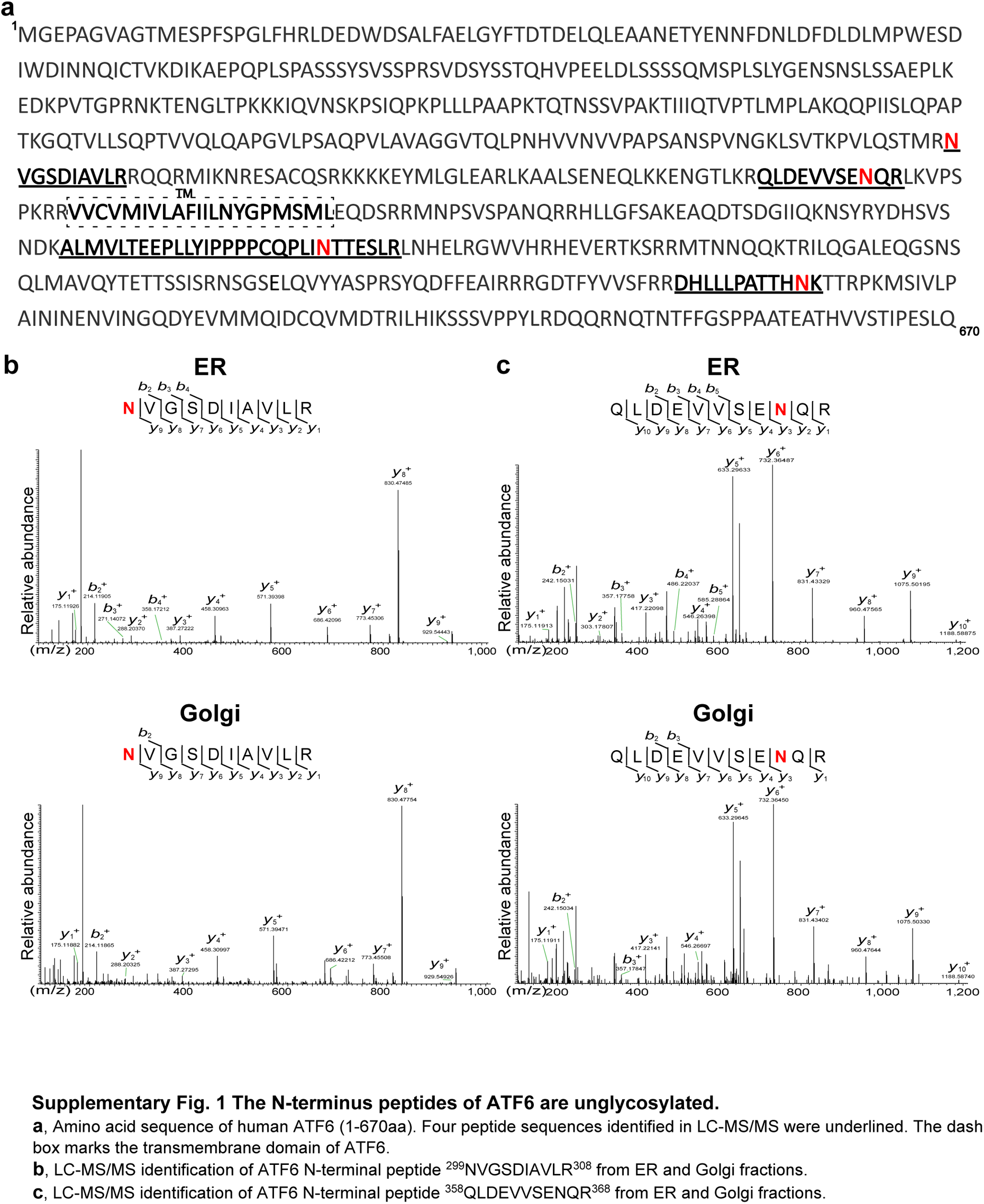
The N-terminus peptides of ATF6 are unglycosylated. **a**, Amino acid sequence of human ATF6 (1-670aa). Four peptide sequences identified in LC-MS/MS were underlined. The dash box marks the transmembrane domain of ATF6. **b**, LC-MS/MS identification of ATF6 N-terminal peptide ^299^NVGSDIAVLR^308^ from ER and Golgi fractions. **c**, LC-MS/MS identification of ATF6 N-terminal peptide ^358^QLDEVVSENQR^368^ from ER and Golgi fractions.

**Supplementary Fig. 2.**
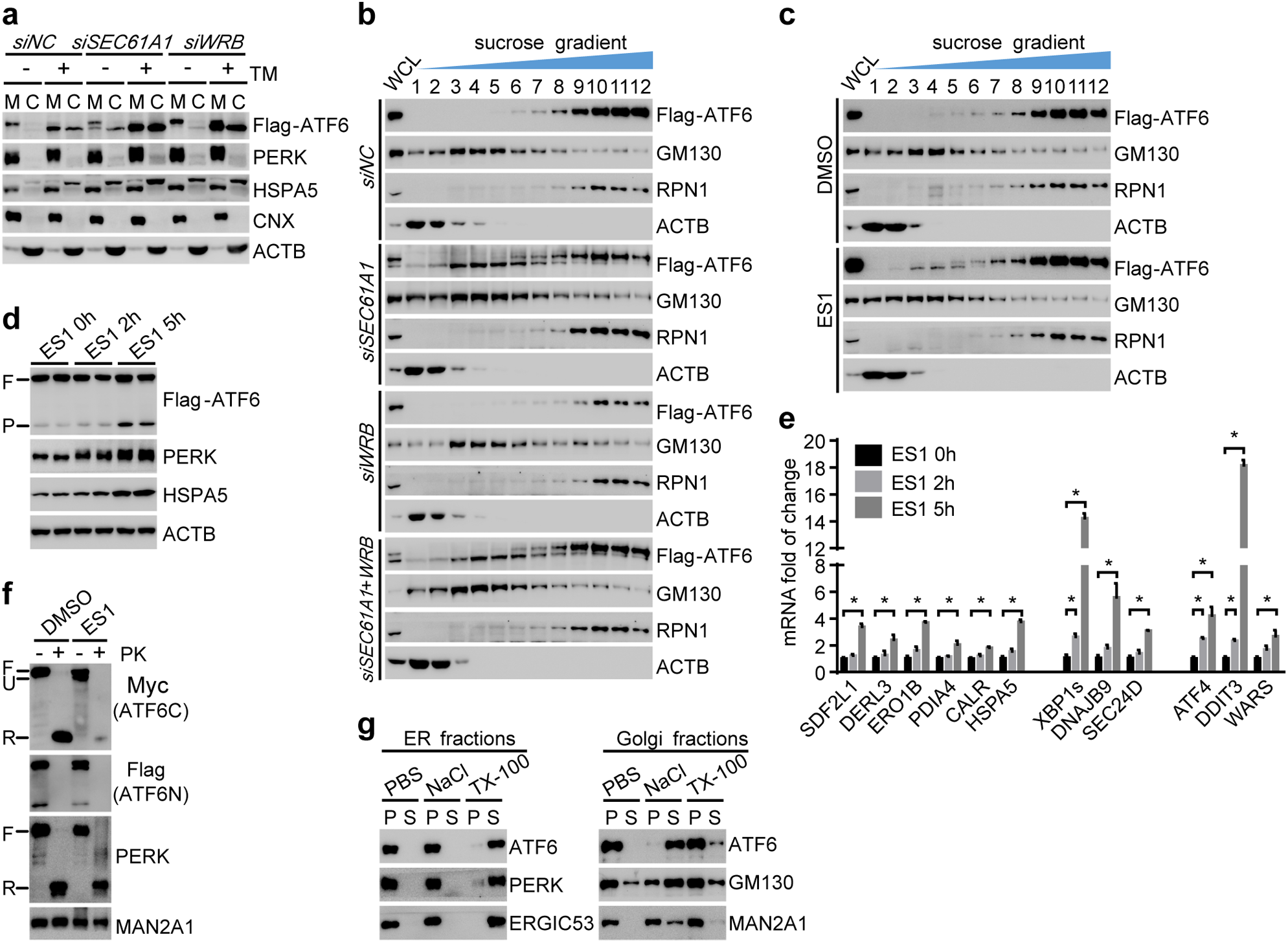
Translocon blockage promotes ATF6 trafficking to the Golgi. **a**, IB of ATF6 in crude fractionation. Cells were transfected with scramble or target gene siRNA and collected with or without ER stress induction (5ug/ml Tm, 5h). M, heavy membrane; C, light membrane and cytosol. **b**, A complete version of **Fig. 3b**. **c**, IB of ATF6 across discontinuous sucrose gradient fractions. Cell culture samples were collected with or without translocon blocker treatment (20nM ES1, 5h). **d**, IB of ATF6 processing during ES1 (20nM) treatment. **e**, qPCR measurement of the induction of UPR genes by ES1 (20nM) treatment. Data are means ± SEM, n=4, *p < 0.05, multiple t-tests. **f**, Proteinase K protection assay of ATF6. Cell culture samples were collected with or without ES1 (20nM, 5h) treatment. **g**, Topology analysis of ER- and Golgi-bound ATF6 from (**c**) with salt wash (0.5 M NaCl) and detergent treatment (2% TX-100). ATF6 was detected by Flag Ab in ER fractions and by Myc Ab in Golgi fractions.

**Supplementary Fig. 3.**
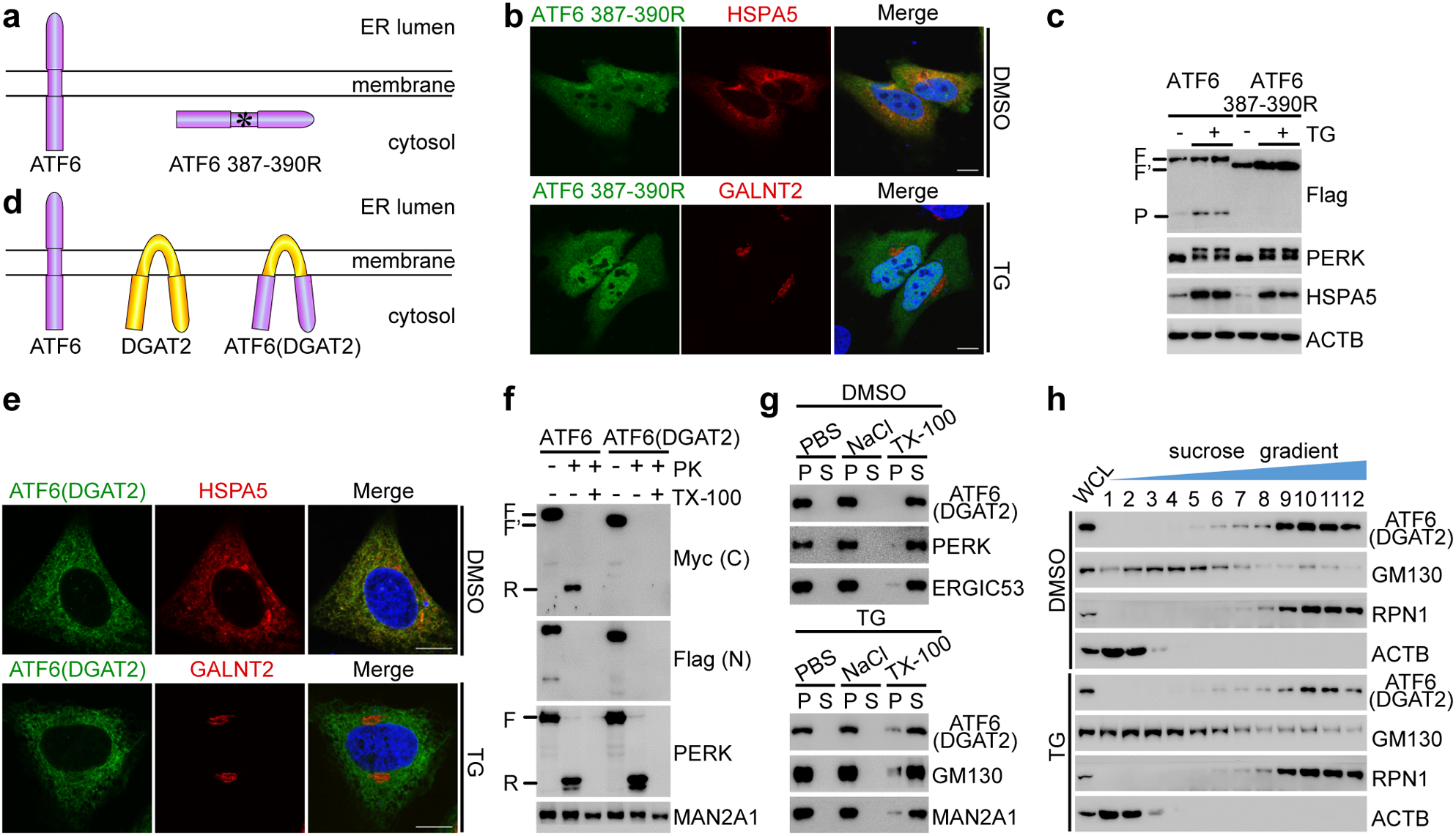
The transmembrane domain of ATF6 is critical for its membrane targeting. **a**, Illustration of the topologies of the wild-type and the mutant ATF6 (FILL387-390RRRR). **b**, IF of ATF6 mutant (387-390R). Cells were fixed with or without TG treatment (100nM, 1h). ATF6 mutant was detected by the N-terminal Flag tag. Scale bar, 10 um. **c**, IB of ATF6 387-390R processing. Cell culture samples were collected with or without TG treatment (100nM, 5h). “F”, full-length of ATF6; “F’”, full-length of ATF6 387-390R. “P”, processed ATF6. **d**, Illustration of the topologies of the wild-type ATF6 (purple), DGAT2 (yellow), and the ATF6(DGAT2) chimera protein on the ER membrane. **e**, IF of ATF6(DGAT2) in normal and stressed (100nM TG, 1 h) cells. ATF6(DGAT2) was detected by the N-terminal Flag tag. Scale bar, 10 um. **f**, Proteinase K protection assay of wild-type ATF6 and ATF6(DGAT2) chimera. “F”, full-length of ATF6; “F’”, full-length of ATF6(DGAT2). “R”, the remaining fragment protected in the ER lumen. **g**, Topology analysis of ATF6(DGAT2) (Flag) with salt wash (0.5 M NaCl) and detergent treatment (2% TX-100). Cell culture samples were collected with or without prior ER stress induction (100nM TG, 5 h). **h**, IB of ATF6(DGAT2) (Flag) across sucrose gradient fractions. Cell culture samples were collected with or without prior ER stress induction (100nM TG, 5 h).

**Supplementary Fig. 4.**
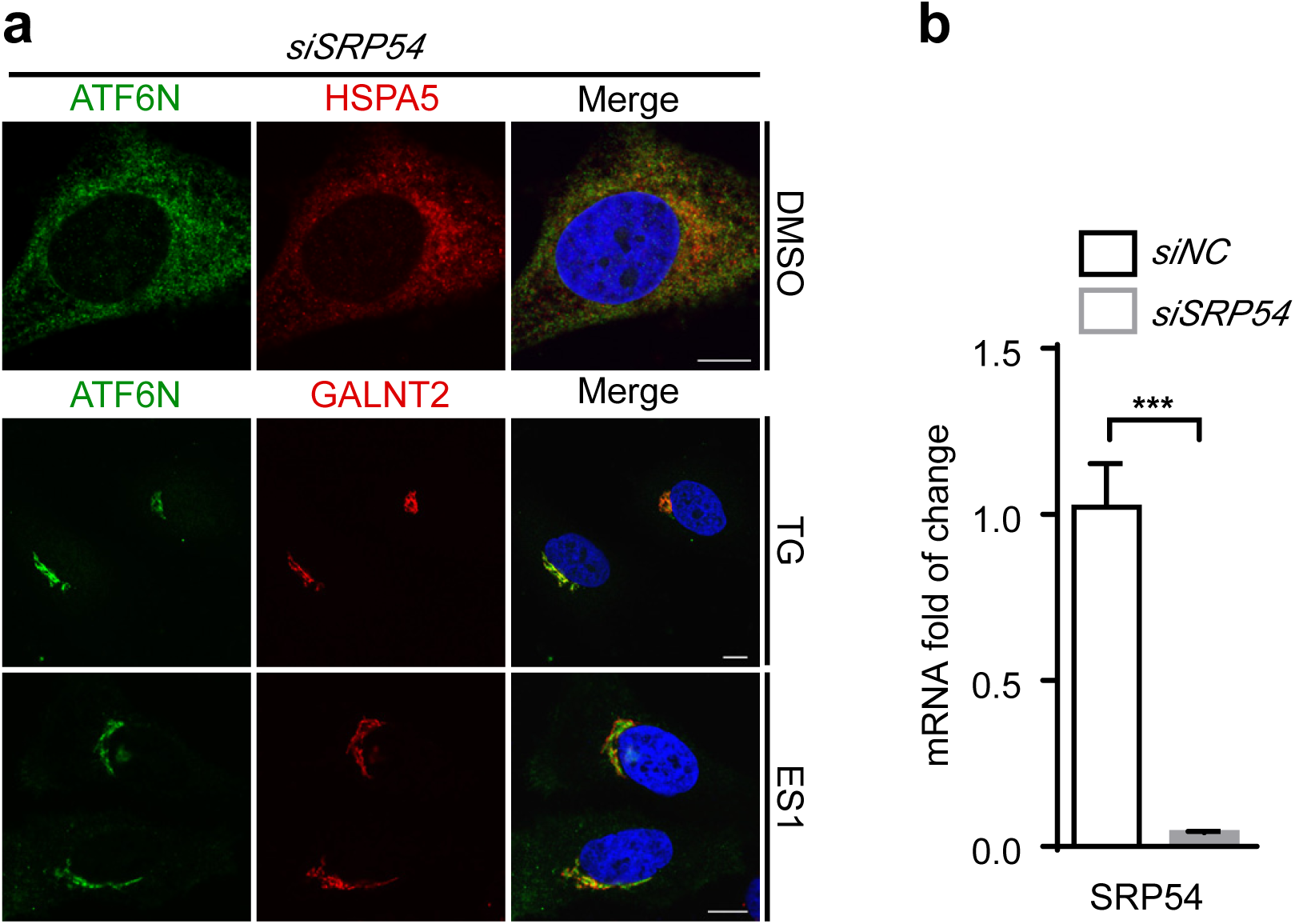
SRP54 knockdown does not affect ATF6 localization. **a**, IF staining of ATF6 (Flag). Cells were transfected with *SRP54* siRNA for 24hs and fixed after either 1h of TG (100nM) or 2hs of ES1 (20nM) treatment. Scale bar, 10 um. **b**, qPCR of SRP54 transcript levels in control and *siSRP54* cell culture samples. Data are means ± SEM, n=4, ***p < 0.001, multiple t-tests.

**Supplementary Fig. 5.**
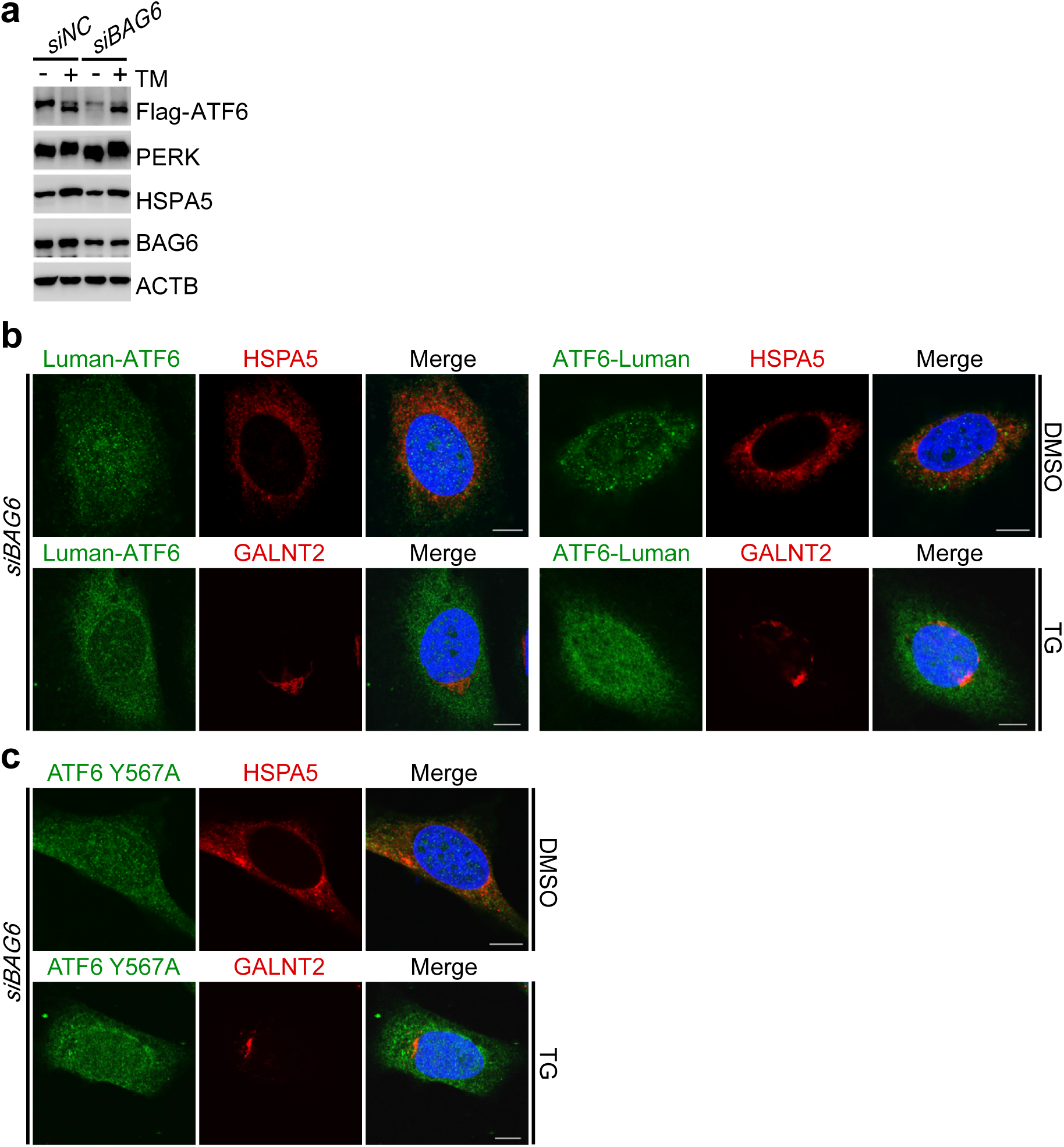
The luminal domain of ATF6 interferes BAG6-dependent membrane insertion under stress conditions. **a**, IB of ATF6. Cells were transfected with indicated siRNA for 24hs were collected with or without prior ER stress inductions (5ug/ml TM, 5h). *siBAG6* contained siRNAs targeting three components of the BAG6 complex: *BAG6, GET4*, and *UBL4A*. **b**, IF of ATF6 chimera proteins (Flag). Cells were transfected with *BAG6* siRNA for 24hs were fixed with or without prior TG treatment (100nM, 1h), compared to **Fig. 4b**. Scale bar, 10 um. **c**, IF staining of ATF6 Y567A (Flag), compared to **Fig. 4g**. Scale bar, 10 um.

**Supplementary Fig. 6.**
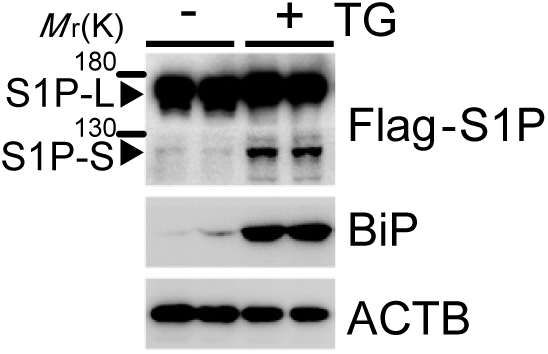
ER stress induces the appearance of a fast-migrating S1P. IB of S1P. Total lysates were prepared from control and TG-treated (100nM, 10 h) cell culture samples. “L”, “S”, different molecular weight of S1P.

**Supplementary Fig. 7.**
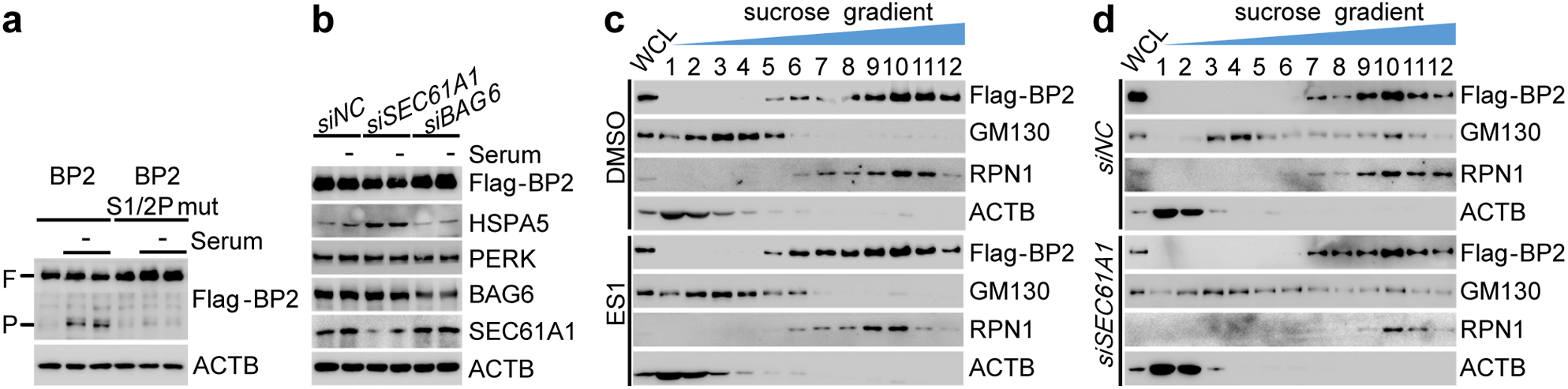
Translocon blockage accelerates SREBP2 activation after serum withdrawal. **a**, IB of wild-type and mutant SREBP2. Samples were subject to 5hs of serum withdrawal before collected. “-”, serum starvation. **b**, IB of SREBP2. Cells were transfected with siRNAs targeting indicated genes for 24hs and collected with or without subject to 5hs of serum withdrawal. **c**, IB of SREBP2 across sucrose gradient fractions. Cell culture samples were collected with or without prior ES1 treatment (20nM, 2h). **d**, IB of SREBP2 across sucrose gradient fractions from control and *siSEC61A1* cells.

