## Supplementary material for "ER stress drives ER-to-Golgi trafficking of ATF6 by blocking its membrane insertion": method

### Materials and methods

#### Key resources table

| Designation | Source | Identifiers |
|---|---|---|
| **Antibodies** | | |
| Mouse monoclonal anti-DYKDDDDK-tag (3B9) | Abmart | Cat#M20008L |
| Mouse monoclonal anti-Myc-tag (9B11) | Cell Signaling | Cat#2276S; RRID:AB_331783 |
| Rabbit polyclonal anti-GFP | Bioeasy | Cat#BE2002 |
| Rabbit polyclonal anti-BiP | Protein tech | Cat#11587-1-AP |
| Rabbit monoclonal anti-PERK (D11A8) | Cell Signaling | Cat#5683; RRID:AB_10841299 |
| Mouse monoclonal anti-MAN2A1(D-5) | Santa Cruz | Cat#sc-377204 |
| Rabbit polyclonal anti-GM130 | Protein tech | Cat#11308-1-AP |
| Rabbit polyclonal anti-RPN1 | Protein tech | Cat#12894-1-AP |
| Mouse monoclonal anti-Beta-Actin | Abgent | Cat#AM1021B |
| Rabbit polyclonal anti-ERGIC-53 | Protein tech | Cat#13364-1-AP |
| Rabbit polyclonal anti-Calnexin | Ayabio | Cat#AYA34411 |
| Rabbit polyclonal anti-WRB | Life Technologies | Cat#PA5-76206 |
| Rabbit monoclonal anti-SEC61A1 (D4K2Z) | Cell Signaling | Cat#14867 |
| Rabbit polyclonal anti-BAG6 | Protein tech | Cat#26417-1-AP |
| Rabbit polyclonal anti-ASNA1 | Protein tech | Cat#15450-1-AP |
| Rabbit polyclonal anti-GALNT2 | Thermo Fisher | Cat#PA5-21541; RRID:AB_11152749 |
| Goat anti-mouse IgG (H+L), HRP conjugated | Bioeasy | Cat#BE0102 |
| Goat anti-rabbit IgG (H+L), HRP conjugated | Bioeasy | Cat#BE0101 |
| Mouse anti-rabbit IgG (Light chain), HRP conjugated | Bioeasy | Cat#BE0107 |
| Goat-anti mouse lgG (H+L)-Alexa Fluor 488 | Life Technologies | Cat#A11001 |
| Goat anti-rabbit lgG (H+L)-Alexa Fluor 568 | Life Technologies | Cat#A11036 |
| **Chemicals, Peptides, and Recombinant Proteins** | | |
| Thapsigargin | Sigma-Aldrich | Cat#T9033 |
| Tunicamycin | Abcam | Cat#ab120296 |
| $H_2{}^{18}O$ | Cambridge Isotope Laboratories (CIL) | Cat#W-0434115 |
| Eeyarestatin 1 | R&D | Cat#3922 |
| Digitonin | Sigma-Aldrich | Cat#11024-24-1 |
| Proteinase K | NEB | Cat#P8107 |
| **Critical Commercial Assays** | | |
| FastQuant Reverse Transcription Kit | Tiangen | Cat#KR116-02 |
| BCA protein assay kit | Solarbio | Cat#PC0021 |
| One step cloning kit | Vazyme | Cat#C112-01 |
| **Experimental Models: Cell Lines** | | |
| HeLa | ATCC | CCL-2 |
| HEK293 | ATCC | CRL-1573 |
| HeLa Flag-ATF6-MYC | This paper | N/A |
| HeLa Flag-ATF6-MYC S1/2m | This paper | N/A |
| HeLa Flag-ATF6(DGAT2)-MYC S1m | This paper | N/A |
| HeLa Flag-LUMAN-ATF6-MYC S1/2m | This paper | N/A |
| HeLa Flag- ATF6-LUMAN-MYC S1/2m | This paper | N/A |
| HeLa Flag-ATF6 Y567A-MYC | This paper | N/A |
| HeLa Flag-ATF6 Y567A-MYC S1/2m | This paper | N/A |
| HeLa MYC(453aa)-S1P-GFP-Flag | This paper | N/A |
| HeLa Flag-SREBP2 | This paper | N/A |
| HeLa Flag-SREBP2 S1/2m | This paper | N/A |
| HEK293 Flag-ATF6-MYC | This paper | N/A |

*Continued*

| Designation | Source | Identifiers |
|---|---|---|
| HEK293 Flag-ATF6-MYC S1/2m | This paper | N/A |
| HEK293 Flag-ATF6(DGAT2)-MYC S1m | This paper | N/A |
| HEK293 Flag-LUMAN-ATF6-MYC S1/2m | This paper | N/A |
| HEK293 Flag- ATF6-LUMAN-MYC S1/2m | This paper | N/A |
| HEK293 Flag-ATF6 Y567A-MYC | This paper | N/A |
| HEK293 Flag-ATF6 Y567A-MYC S1/2m | This paper | N/A |
| HEK293 MYC(453aa)-S1P-GFP-Flag | This paper | N/A |
| HEK293 Flag-SREBP2 | This paper | N/A |
| HEK293 Flag-SREBP2 S1/2m | This paper | N/A |
| **Recombinant DNA** | | |
| Flag-ATF6-Myc | This paper | N/A |
| Flag-ATF6-Myc N391F P394L R416A L419V | This paper | N/A |
| Flag-ATF6(1-377)-DGAT2(90-112)-ATF6(399-670)-Myc | This paper | N/A |
| Flag-ATF6(1-377)-DGAT2(90-112)-ATF6(399-670)-Myc R416A L419V | This paper | N/A |
| Flag-Luman(1-229)-ATF6(378-670)-Myc | This paper | N/A |
| Flag-Luman(1-229)-ATF6(378-670)-Myc N391F P394L R416A L419V | This paper | N/A |
| Flag-ATF6(1-430)-Luman(270-371)-Myc | This paper | N/A |
| Flag-ATF6(1-430)-Luman(270-371)-Myc N391F P394L R416A L419V | This paper | N/A |
| Flag-ATF6 Y567A-Myc | This paper | N/A |
| Flag-ATF6 Y567A-Myc N391F P394L R416A L419V | This paper | N/A |
| Flag-ATF6-Myc 387-390RRRR | This paper | N/A |
| Flag-ATF6-Myc 387-390RRRR N391F P394L R416A L419V | This paper | N/A |
| Flag-ATF6 | This paper | N/A |
| Flag-ATF6 387-390RRRR | This paper | N/A |
| Flag-ATF6 Y567A | This paper | N/A |
| Flag-Luman(1-229)-ATF6(378-670) | This paper | N/A |
| Flag-ATF6(1-430)-Luman(270-371) | This paper | N/A |
| MYC(453aa)-S1P-GFP-Flag | This paper | N/A |
| Flag-SREBP2 | This paper | N/A |
| Flag-SREBP2 R519A L522V N495F P496L | This paper | N/A |
| **Sequence-Based Reagents** | | |
| siRNA targeting sequence: SEC61A1: CCAACCUCAUGAAUCUCAUTT | This paper | N/A |
| siRNA targeting sequence: WRB: GCUGUCGUGCCGAGUAAAUTT | This paper | N/A |
| siRNA targeting sequence: SRP54: GCGAGACAUGUAUGAGCAATT | This paper | N/A |
| siRNA targeting sequence: BAG6: CCAGAUGGUGAGCGGCCUUTT | This paper | N/A |
| siRNA targeting sequence: GET4: GCGUCGAGAAGGGCGACUATT | This paper | N/A |
| siRNA targeting sequence: UBL4A: GCUCAACCUAGUGGUCAAATT | This paper | N/A |
| QPCR primers see table S2 | This paper | N/A |
| Clone primers see table S2 | This paper | N/A |
| **Software and Algorithms** | | |
| Illustrator CC 2015 | http://www.adobe.com/products/illustrator.html | N/A |

*Continued*

| Designation | Source | Identifiers |
|---|---|---|
| Photoshop CS6 | http://www.adobe.com/cn/products/photoshop.html | N/A |
| AlphaView (FluroChem FC3) | ProteinSimple | N/A |
| Prism 6 | http://www.graphpad.com/scientific-software/prism/ | N/A |
| Volocity | PerkinElmer | N/A |
| Proteome Discovery searching algorithm (1.4) | Thermo | N/A |
| **Others** | | |
| Anti-FLAG® M2 Magnetic Beads | Sigma-Aldrich | Cat#M8823-1ML |
| Protein A/G Agarose beads | CMCTAG | Cat#IF0001 |
| TRIzol Reagent | Invitrogen | Cat#15596026 |
| RNAiMAX | Thermo Fisher | Cat#13778030 |
| TRITON® X-100 | Ameresco | Cat#0694 |

##### Cell Culture

HeLa cells or HEK293 cells were obtained from the American Type Culture Collection, verified by STR testing, and regularly tested to be mycoplasma-free throughout this study. All cells were cultured in DMEM (containing 100 units/ml penicillin and 100 mg/ml streptomycin sulfate) supplemented with 10% fetal bovine serum (FBS) and grown in a monolayer at 37 °C in 5% $CO_2$. Plasmid DNA and small interference RNA were transfected with either polyethyleneimine (Sigma) or Lipofectamine RNAiMAX (Thermo Fisher) according to the manufacturer recommendations.

##### Construction of Plasmids and Stable Cell Lines

Human ATF6, SREBP2, S1P, DGAT2, and Luman were PCR amplified from HEK293 cDNA and cloned into a pEGFP backbone expression vector with a CMV promoter and G418 selection marker. The GFP tag was removed and Flag and Myc tags were added to the 5'- and 3'-end of the multiple cloning site (MCS). Site-directed mutagenesis is carried out with the QuikChange® II XL Site-Directed Mutagenesis Kit (Agilent) according to manufacturer's recommendations. For chimera construction, fragments of ATF6, DGAT2, and Luman were PCR-amplified and sub-cloned with the One Step Cloning kit (Vazyme) according to manufacturer's recommendations. For the construction of stable cell lines, HeLa cells or HEK293 cells were transfected with appropriate plasmids, and single cell colonies were selected by antibiotics, screened for their expression of desired proteins, and expanded for future experiments. All primers used for plasmid construction were listed in STAR table.

##### Mass spectrometry (MS) analysis of glycosylation peptides by $^{18}$O labeling

Protein lysates from ER and Golgi fractions were subjected to SDS-PAGE and stained with Coomassie Blue. Gel slices from the expected size of ATF6 were de-stained, reduced with 25 mM DTT at 57 °C for 30 min, and alkylated with 50 mM iodoacetamide at room temperature for 20 min. Then the gel pieces were incubated with PNGase F (500 U) in the presence of $^{18}$O-water with 95% isotope purity at 37 °C for 24 hours. After incubation, in-gel digestion was performed using sequencing grade-modified trypsin (1.5 ug) in 50 mM ammonium bicarbonate at 37 °C overnight. The peptides were extracted with of 50% ACN aqueous solution containing 0.1% TFA. For LC-MS/MS analysis, peptides were separated by a 120 min gradient elution at a flow rate 0.300 μL/min with the EASY-nLC 1000 system which was directly interfaced with the Thermo Orbitrap Fusion mass spectrometer.

##### Immunofluorescence and Cell Imaging

HeLa cell stable lines expressing designated constructs were plated on glass slips for overnight growth before chemical treatments. Cells were then washed in phosphate-buffered saline (PBS), fixed in 4% paraformaldehyde for 10 min at room temperature, permeabilized and blocked with 0.7% Triton X-100/5% FBS for 1 hour, stained with Flag (or Myc) and HSPA5 (or GALNT2) diluted 1:200 in PBS for 1 hour at room temperature. Cells were then washed three times in PBS, and incubated with Alexa488 goat anti-mouse (for Flag or Myc) and Alexa568 goat anti-rabbit (for HSPA5 or GALNT2) secondary antibodies at 1:1000 dilutions in PBS for 1 hour at room temperature. Cells were washed for three more times in PBS and stained with Hoechst33342 (Invitrogen) at 1:1000 dilutions for 10 min at room temperature. Cells were washed again for three more times in PBS and mounted in Vectashield mounting medium H1000 (Vector Laboratories) before imaging with FV1200 confocal microscope.

For each co-localization analysis, 100 cells from two independent experiments were imaged by UltraVIEW Vox spinning disk confocal microscope through Z-axis stacking, analyzed in Volocity 3D Image Analysis Software (PerkinElmer), and the Pearson's correlation coefficient was calculated.

**Fluorescence Proteinase K Protection (FPP) assay**

For fluorescence proteinase K protection assay, cells were plated on poly-d-lysine (Beyotime) coated glass slips. After an overnight culture, cells were washed in PBS, permeabilized with 12 uM digitonin (Sigma), and digested by 100 ug/ml proteinase K for 5 min in HBSS supplied with calcium and magnesium (Gibco). Cells were then washed, fixed, and prepared for immunofluorescence imaging as described above.

**Proteinase K Protection Assay**

HEK293 stable lines expressing desired proteins were plated in a 6cm dish and treated with indicated drugs before sample collection. Cells were then lysed in 500 ul ice-cold homogenization buffer containing 10 mM HEPES-KOH (PH 7.5), 220 mM D-mannitol and 70 mM sucrose, and homogenized with 30 strokes using 29-gauge needles. Cell lysates were centrifuged at 1000 g for 10 min at 4 °C to remove intact cells and nuclei (debris). The supernatant was centrifuged at 100,000 g for 1 hour at 4 °C in an Optima MAX Ultracentrifuge (Beckman Coulter, TLA 55). The resulting pellet was resuspended in homogenization buffer and aliquoted. For proteinase K protection assay, 50 μg/mL proteinase K (NEB) with or without 0.5% TritonX-100 were added to each aliquot and incubated at 4 °C for 30 min. Proteins were precipitated with 10% trichloroacetic acid on ice for 30 min, then centrifuged at 12,000 g for 20 min at 4 °C. Supernatants were discarded, and pellets were washed twice with ice-cold acetone. The pellets were air dried and solubilized in 1x loading buffer with 3 M urea and heated at 95 °C for 5 min before loading onto SDS-PAGE.

**Discontinuous Sucrose Gradient Fractionation**

HEK293 stable lines expressing desired protein were plated in a 10cm dish and treated with indicated drugs before sample collection. Cells were then collected and lysed in 1ml ice-cold sucrose buffer containing 10 mM HEPES-KOH (pH 7.4), 0.25 M sucrose, 1 mM EDTA, protease inhibitor cocktail (Selleck), and homogenized with 50 strokes using 29-gauge needles. Cell lysates were centrifuged at 1000 g for 10 min at 4°C to remove intact cells and nuclei (debris). Two hundred microliter aliquots of the resulting supernatant were laid on top of a discontinuous sucrose gradient (20%, 30%, 40%, 50%, and 60% from top to bottom, 400ul each), centrifuged at 4°C at 50,000 rpm for 30 min (TLS 55, Optima MAX Ultracentrifuge, Beckman Coulter), and stopped without braking. Twelve fractions of 100ul each were collected from the top and used for downstream analyses.
For immunoblot analysis, the aliquots were precipitated with 10% trichloroacetic acid on ice for 30min followed by centrifugation at 12,000 g for 20 min at 4°C. For proteinase K protection assay, each fraction was diluted to a final concentration of 10% sucrose, pelleted at 100,000 g for 3 hours at 4 °C (TLA 55, Optima MAX Ultracentrifuge, Beckman Coulter), washed and resuspended in homogenization buffer. For salt wash and detergent treatments, each fraction was diluted to 10% sucrose supplemented with 5 mM MgCl2 and 50 mM NaCl, centrifuged at 100,000 g for 3 hours at 4°C, resuspended in sucrose buffer, and treated with either 500 mM NaCl, 2% TritonX-100, or PBS control for 15 min at room temperature. The suspension was then centrifuged at 100,000 g for one hour to obtain the supernatant and a membrane pellet.

**Crude Fractionation**

HEK293 stable lines expressing designated proteins were plated in a 6cm dish and transfected with indicated siRNAs for 24 hours and treated with indicated drugs. Cells were then collected and swelled at 0 °C for 30 min in a 500 ul buffer B containing 10 mM HEPES-KOH (pH7.4), 10 mM KCl, 1.5 mM $MgCl_2$, 0.5 mM sodium EDTA (pH8.0), 0.5 mM sodium EGTA (pH8.0), 1 mM DTT, protease inhibitor cocktail (Selleck). The lysate was cleared by centrifugation at 1000 g for 10 min at 4 °C to remove debris and the resultant supernatant was centrifuged at 100,000 g for 20 min to obtain a microsomal pellet and a cytosolic supernatant.

**Immunoblot**

Whole cell lysates were prepared in a RIPA lysis buffer (20 mM Tris-HCl, pH7.5, 150 mM NaCl, 1 mM EDTA, 1 mM EGTA, 1% NP-40, 1% sodium deoxycholate) supplemented with protease inhibitor cocktail (Selleck). Protein concentrations were quantified by BCA kit (Solarbio) and denatured at 95 °C for 5 min. A total of 25 ug proteins were subjected to SDS-PAGE and blotted onto nitrocellulose membrane (Pall Corporation), blocked with 5% milk (BD Biosciences), incubated with the indicated primary antibodies and corresponding HRP-conjugated secondary antibodies, developed with ECL reagents (Pierce), visualized and analyzed by AlphaView software (ProteinSimple).

**Immunoprecipitation**

Cells were transfected for 24 hours with designated expression plasmids prior to drug treatment. Cell lysates were then prepared in 0.3% TritonX-100 lysis buffer (20 mM HEPES-NaOH, pH 7.5, 150 mM NaCl, 10% glycerol, 1 mM EDTA) supplemented with protease inhibitor cocktail (Selleck), and cleared at 1000 g for 10 min at 4 °C. For BAG6 immunoprecipitation, crude lysates were incubated with BAG6 or IgG antibody for 2 hours at 4°C, followed by the addition of protein-A/G-Agarose Beads (CMCTAG) for 90 min at 4 °C, pelleted by 100 g for 1 min. For Flag immunoprecipitation, crude lysates were incubated with Flag-conjugated beads (Sigma) for 2 hours and magnetically captured. Beads were washed three times with 1 ml of lysis buffer and resuspended in 1X protein sample buffer for SDS-PAGE.

**Reverse Transcribed Quantitative Polymerase Chain Reaction (RT-qPCR)**

Total RNA was isolated using TRIzol reagent (Invitrogen) and reverse transcribed with the FastQuant RT Kit (Tiangen). cDNA was quantified by real-time PCR using SYBR Green Master Mix (Thermo Fisher) on an ABI 7900 (ThermoFisher). Duplicate runs of each sample were normalized to GAPDH or 18s to determine relative gene expression levels. Values plotted to represent averages from at least four different biological samples. Primer sequences were listed in the STAR table.

**Statistical analysis**

The number of replicates were described in the figure legends. For biochemical assays, n corresponds to the number of experimental replicates. For imaging analyses, n corresponds to the number of cells. Immunoblot quantification was performed by AlphaView (ProteinSimple). Co-localization quantification was performed in Volocity 3D Image Analysis Software (PerkinElmer). All data are presented as means ± SEM and analyzed using GraphPad Prism6. Student's t-test was used for single variable comparison between two groups. Two-way ANOVA was used to examine interactions between multiple variables. *p*-value < 0.05 was considered to be statistically significant and is presented as * $p < 0.05$, ** $p < 0.01$, *** $p < 0.001$.
